# A Transient Immunostimulatory Niche Synergizes Adoptive and Endogenous Immunity for Enhanced Tumor Control

**DOI:** 10.1101/2025.08.22.671875

**Authors:** Anahita Nejatfard, John H. Klich, Noah Eckman, Sophia J. Bailey, Kyra Gillard, Daniel Ramos Mejia, Ben S. Ou, Jerry Yan, John W. Hickey, Eric A. Appel

## Abstract

Adoptive Cell Therapy (ACT) has achieved curative responses in hematological malignancies, yet its translation to solid tumors remains limited by manufacturing bottlenecks, systemic toxicities, and poor T-cell infiltration and persistence within the immunosuppressive tumor microenvironment (TME). Here, we report the development and mechanism of ACTIVATE (Adoptive Cell Therapy and Immunostimulatory Vehicle for Anti-Tumor Efficacy), which leverages an injectable hydrogel depot technology that forms a transient inflammatory niche for localized co-delivery of adoptive T cells and native cytokines. By tuning cytokine identity, ACTIVATE enables precise modulation of T-cell expansion, effector function, and interaction with endogenous immune networks. We found that enhancing T-cell proliferation alone is insufficient to drive robust tumor control; instead, coordinated engagement of both adoptive and endogenous immune responses is critical for durable anti-tumor efficacy. *In vivo*, this orchestration via ACTIVATE led to enhanced infiltration and cytotoxicity of both adoptive and host-derived immune effectors, while driving robust recruitment and activation of T cells, B cells, dendritic cells, and macrophages in the tumor-draining lymph nodes. This local immune activation can further reshape the TME, promoting antigen presentation and suppressing immunoregulatory populations, thus enhancing anti-tumor efficacy in murine melanoma and lymphoma models. These findings establish ACTIVATE as a modular platform for orchestrating coordinated immune responses to improve ACT outcomes in solid tumors.

## 1. Introduction

Cancer remains a leading cause of mortality worldwide, with an estimated two million new cases projected in the United States in 2025^1^. Adoptive cell therapy (ACT), a revolutionary immunotherapy involving the genetic engineering of T cells to express chimeric antigen receptors (CARs) or modified T-cell receptors (TCRs) for enhanced tumor recognition, has demonstrated unprecedented efficacy in advanced hematologic malignancies^2^. To date, six CAR-T cell therapies have received U.S. Food and Drug Administration approval, with numerous ongoing clinical trials expanding the therapeutic landscape^3^. Yet, widespread implementation remains constrained by the necessity for extensive *ex vivo* expansion, a laborious and costly process that can impair T-cell fitness upon reinfusion^4,5^.

Beyond manufacturing bottlenecks, systemic ACT administration poses significant safety concerns. On-target off-tumor toxicity (OTOT) arises when adoptively transferred cells target antigens expressed on normal tissues, leading to collateral damage^6^. Additionally, cytokine release syndrome (CRS), a severe inflammatory response driven by excessive immune activation, necessitates prophylactic administration of immunosuppressive agents like tocilizumab, an IL-6 inhibitor, in over 70% of patients receiving FDA-approved CAR-T therapies^7,8^. Further, translating the efficacy of ACT to solid tumors remains elusive. Intravenously administered adoptive cells face formidable challenges in trafficking to and infiltrating the tumor site^9^. Even when tumor infiltration occurs, the immunosuppressive tumor microenvironment (TME) hinders T-cell expansion and function, often leading to T-cell exhaustion, a dysfunctional state characterized by reduced cytotoxicity and impaired persistence^9,10^.

To circumvent these barriers, localized ACT delivery has emerged as a promising strategy to enhance intratumoral T cell accumulation while mitigating systemic toxicity, with early preclinical results demonstrating improved outcomes^11,12^. Unfortunately, overcoming both T cell exhaustion and the immunosuppressive TME remain major obstacles. Cytokines—key modulators of immune activation, expansion, and polarization—play a vital role in ACT manufacturing and can enhance *in vivo* persistence and functionality of adoptively transferred cells^13–15^. Among cytokines, interleukins are highly significant to the development and progression of cancer, with active roles in tumor progression and control^16^. Yet, the systemic cytokine concentrations necessary to achieve desired effects often result in severe toxicities, limiting their clinical application outside of desperate cases^13,17^. Locoregional cytokine delivery offers a means to sustain T-cell activity while minimizing adverse systemic effects^18^.

In the interest of improving locoregional ACT delivery, biomaterial-based platforms incorporating stimulatory cytokines have been explored to promote T-cell expansion, persistence, and infiltration^19–21^. While these approaches show promise, current biomaterial platforms face translational challenges, including limited scalability, restricted tunability for diverse tumor types, and complex clinical implementation^7,21,22^. Many require invasive implantation or have been validated only in immune-privileged tissue and immunocompromised murine models^7,22^. Additionally, most existing biomaterial platforms necessitate engineering cytokines into specialized formulations—such as microspheres or insoluble depots—to prevent rapid diffusion away from the tumor site, limiting flexibility in cytokine selection^20,23,24^. Moreover, these approaches primarily focus on adoptive cell persistence without adequately engaging endogenous immunity to generate a more robust and durable anti-tumor response. This disparity highlights the need for next-generation ACT delivery platforms that not only support adoptive T-cell persistence and function but also harness the broader immune system for long-term tumor control.

To address these limitations, we previously engineered an injectable polymer-nanoparticle (PNP) hydrogel depot technology that uniquely enables prolonged retention of interleukin-15 (IL-15) while supporting the sustained expansion and enhanced tumor reactivity of encapsulated CAR-T cells^25^. Preliminary studies validated the therapeutic potential of this approach; however, the precise mechanisms driving CAR-T cell activity within this system remain poorly understood, largely due to the reliance on xenogeneic CAR-T models^26^. It remains unclear, for instance, whether the observed enhancement in tumor reactivity is directly attributable to cytokine co-delivery or if alternative mechanisms contribute to this effect. Whether the prolonged retention observed for IL-15 in these PNP hydrogels is unique to this cytokine or extends to others also remains an open question. Beyond ACT, the dynamic nature of PNP hydrogels has been leveraged in vaccine strategies to recruit immune cells into the depot, amplify immune responses, and shape broader antiviral immunity^27^. However, in the context of ACT, such immune interactions remain underexplored. The use of xenograft models—where human CAR-T cells are evaluated against human tumors in immunocompromised mice—precludes key immune interactions that could potentiate ACT efficacy.

To bridge these gaps, we developed ACTIVATE (Adoptive Cell Therapy and Immunostimulatory Vehicle for Anti-Tumor Efficacy) (**Fig. 1A**). We hypothesized that evaluating this platform in a syngeneic ACT model would enable a more comprehensive mechanistic investigation by preserving immune interactions absent in xenograft systems. Furthermore, we propose that incorporating alternative cytokines within the hydrogel may differentially shape adoptive T-cell phenotype and function, enhancing persistence, memory formation, exhaustion resistance, and synergy with complementary therapies such as immune checkpoint blockade (ICB). Through these investigations, we aim to elucidate the full immunological potential of ACTIVATE and establish a robust framework for its clinical translation.

**Figure 1.**
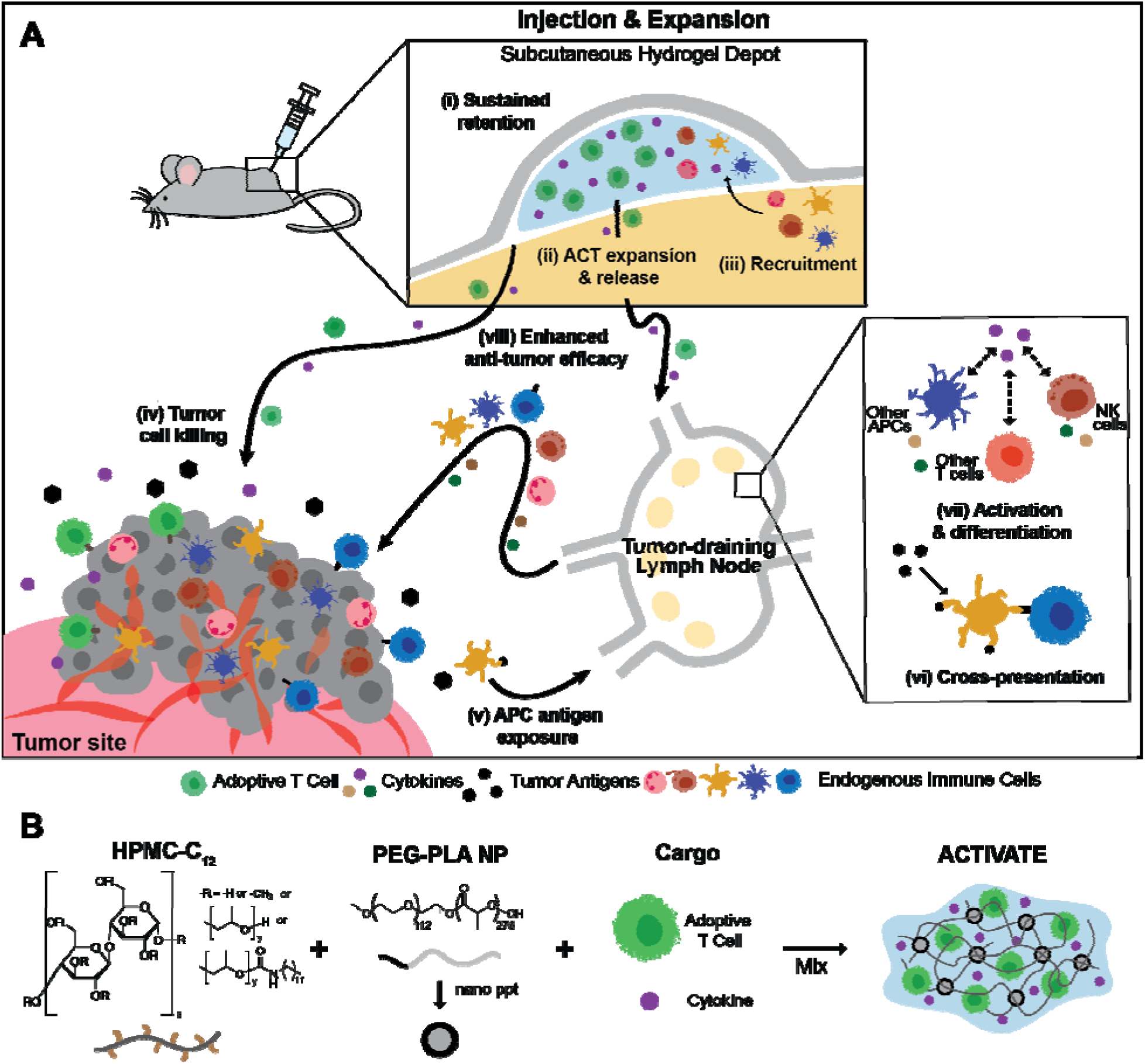
Adoptive Cell Therapy and Immunostimulatory Vehicle for Anti-Tumor Efficacy. (**A**) Schematic illustration demonstrating proposed mechanism of ACTIVATE: (i-ii) Injected hydrogel containing adoptive T cells and cytokine(s) enables long-term local retention of cytokine alongside sustained adoptive T cell expansion and release over the course of 2 weeks; (iii) recruitment of endogenous immune cells, generating a local inflammatory niche for activation and expansion of endogenous immune cells; (iv) adoptive T cell and cytokine release elicit potent tumor cell killing, releasing antigens that can be (v) picked up by antigen-presenting cells and taken back to the tumor-draining lymph nodes for (vi) cross-presentation to endogenous T cells; (vii) combined with the activation and differentiation of endogenous immune cells in the tumor-draining lymph nodes provoked by drainage of adoptive T cells and cytokine enable (viii) engagement of endogenous immune cells for enhanced anti-tumor efficacy. (**B**) Polymer-nanoparticle hydrogels formed through self-assembly of dodecyl-modified hydroxypropyl methylcellulose (HPMC-C_12_) and degradable PEG-PLA core-shell nanoparticles, enabling co-encapsulation of adoptive T cells and stimulatory cytokines.

Using well-characterized preclinical models of ACT involving transgenic PMEL CD8^+^ T cells for B16-F10 melanoma and OT-I CD8^+^ T cells for E.G7-Ova lymphoma, we demonstrate that ACTIVATE serves as a transient inflammatory niche for the local, sustained delivery of adoptive T cells and immunostimulatory cytokines, without requiring modification of the therapeutic cargo. By enabling prolonged cytokine retention within the hydrogel depot, ACTIVATE influences T-cell proliferation, activation, and memory phenotype. *In vivo*, ACTIVATE enhances endogenous immune cell recruitment to the tumor, tumor-draining lymph nodes (tdLNs), and the hydrogel niche itself. The co-encapsulation of cytokines within ACTIVATE promotes the expansion and cytotoxicity of both adoptively transferred and endogenous effector cells, amplifying the immune response. Notably, ACTIVATE synergizes with professional antigen-presenting cells (APCs), fostering a more immunostimulatory TME by reducing immunosuppressive cell populations and mitigating T-cell exhaustion, which in turn allows for synergy with other therapeutics like immune checkpoint blockade (ICB) . Together, these findings highlight ACTIVATE as a promising strategy for improving ACT efficacy against solid tumors by enhancing direct adoptive cell function while engaging the endogenous immune system.

## 2. Results

### 2.1. ACTIVATE facilitates sustained cytokine release and influences T cell dynamics

We previously demonstrated that IL-15 could be seamlessly incorporated into the PNP hydrogel platform by simple mixing, where the dynamic mesh architecture of the hydrogel facilitated prolonged cytokine retention despite the small size of the cargo^25^. Given its pivotal role in promoting T-cell proliferation and differentiation into effector subsets^28^, IL-15 incorporation enabled the PNP hydrogel to function as a sustained reservoir for T-cell expansion. Notably, this system enhanced the tumor-reactivity of CAR-T cells and promoted a stem-cell like memory phenotype of the cells compared to CAR-T cells delivered without cytokine support^25^. Given the versatility of this system, we hypothesized that the hydrogel’s cytokine-retention mechanism would extend to other interleukins, and that cytokine co-encapsulation would differentially influence T-cell proliferation and phenotype in a cytokine-dependent manner.

To test this, we selected interleukins with well-established roles in T-cell expansion and functionality within the context of ACT^13^: IL-15 for its ability to drive activation, proliferation, and survival of T cells^28^; IL-2 for its well-characterized capacity to enhance naïve CD8^+^ T-cell expansion and differentiation into cytotoxic T lymphocytes (CTLs)^29^; IL-7 for its critical role in supporting the survival and maintenance of both naïve and memory T cells^30^; and IL-12 for its potent induction of T-cell proliferation and effector function, particularly in CD4^+^ T cells^31^. To evaluate cytokine-specific effects on ACT outcomes, we utilized the syngeneic PMEL murine model, in which transgenic CD8^+^ T cells specifically recognize the murine homologue of the gp100 antigen expressed on B16-F10 melanoma cells^32^. We prepared ACTIVATE hydrogel systems by physically mixing aqueous solutions of dodecyl-modified hydroxypropyl methylcellulose (HPMC-C_12_) and poly(ethylene glycol)-poly(lactic acid) (PEG-PLA) nanoparticles (NPs) to a final concentration of 1 wt% and 5 wt%, respectively (*n.b.*, denoted PNP-1-5). We then added 2.5μg of each respective cytokine and 2×10^6^ of stimulated and activated PMEL CD8^+^ T cells (**Fig. 1B**). Consistent with prior studies, ACTIVATE exhibited robust solid-like mechanical properties enabling prolonged depot persistence, shear-thinning behavior necessary for injectability through fine-gauge needles, rapid self-healing post-injection to prevent burst release, and excellent biocompatibility (**Fig. S1A-C**)^33,34^.

*In vitro* release studies without co-encapsulated T cells demonstrate that over 85% of all interleukins, independent of their molecular weight, are retained within the PNP-1-5 hydrogel after one week under infinite sink conditions (**Fig. 2A-B**). This feature of the PNP hydrogels enables facile formulation of these various cytokines, as well as cytokine mixtures, without the need for bespoke materials development or modification of the cytokines. To assess the impact of cytokines on T-cell proliferation, we performed cell release assays in which ACTIVATE hydrogels were injected onto transwells with large pore sizes that permit T cell migration into the medium below (**Fig. 2C**). These conditions were compared to a PMEL only control, consisting of a direct injection of the same dose of PMEL cells and IL-2 in PBS without hydrogel to the transwell, to evaluate the effect of encapsulation on cell release dynamics. Each cytokine imparted distinct effects on T-cell proliferation: IL-2 and IL-7 significantly enhanced expansion from ACTIVATE (∼2.5-fold and ∼3.5-fold increases in released cell numbers relative to the loaded dose, respectively), whereas IL-15, and particularly IL-12, did not substantially influence T-cell proliferation (**Fig. 2D-G**).

**Figure 2.**
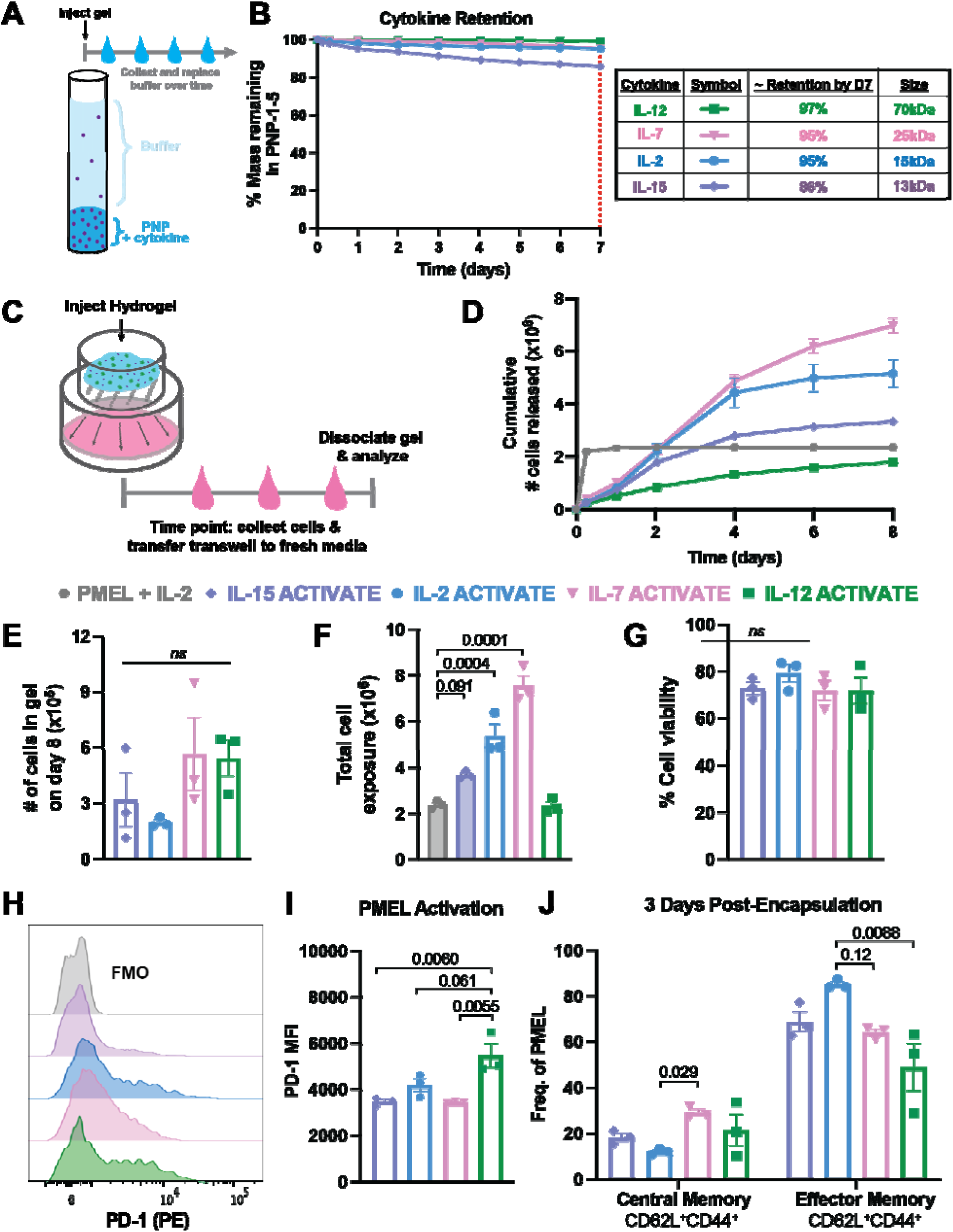
ACTIVATE sustains retention of various cytokines that influence adoptive T-cell expansion and phenotype. (**A**) Schematic of in vitro release assay of cytokines from ACTIVATE immersed in PBS. (**B**) %Mass of cytokines IL-12, IL-7, IL-2, and IL-15 released and remaining in the hydrogels over the 1-week assay. (**C**) Schematic of in vitro cellular release assay: Hydrogel containing PMEL adoptive T cells and cytokine is injected into a porous transwell suspended over medium. Cells proliferate within the hydrogel while also releasing into the medium below. At each timepoint, the medium below the transwell is collected, cells are counted, and transwell is placed into fresh media. (**D**) Cumulative number of cells released into the medium over various timepoints for PMEL and cytokine encapsulated hydrogels and PMEL only control. (**E**) Total number of cells remaining in ACTIVATE on the final day of the assay. (**F**) Total cell exposure determined as the sum of cumulative cells released and total cells remaining in the hydrogel on day 8. (**G**) Cell viability of the cells remaining in the hydrogels on the final day of the assay. (**H**) Representative flow cytometry plots and (**I**) quantification of PD-1 MFI of PMEL cells in the gel 3 days after encapsulation. (**J**) Frequency of central memory (CD62L^+^CD44^+^) and effector memory (CD62L^-^CD44^+^) phenotypes for PMEL cells in the hydrogel 3 days after co-encapsulation with different cytokines. Data are representative of *n* = 3 technical replicates per group and presented as mean ± s.e.m. *P* values were determined by ordinary one-way ANOVA followed by Tukey’s multiple comparison test using GraphPad PRISM.

Most synthetic niche-based systems incorporate integrin-binding motifs to facilitate cell adhesion^35,36^. To assess the role of integrin interaction in our platform, we functionalized the hydrophilic corona of PEG-PLA NPs with the arginine-glycine-aspartic acid (RGD) motif, which we previously demonstrated enhances cell motility and viability^25^. We hypothesized that RGD-functionalization would influence T-cell adhesion and migration within the hydrogel depot. To test this hypothesis, we formulated ACTIVATE with PEG-PLA NPs functionalized with either a low (0.3mM) or high (1mM) concentration of RGD and encapsulated both PMEL CD8^+^ T cells and 2.5μg of IL-2. Quantification of released cells revealed the RGD-functionalized NPs significantly enhances T-cell adhesion, as evidenced by increased retention within the gel eight days post-encapsulation (**Fig. S2A**). Interestingly, while RGD-functionalized hydrogels preserved antigen sensitivity compared to unmodified hydrogels at three days post-encapsulation, this effect was lost by day eight (**Fig. S2B**). Moreover, increased RGD-functionalization significantly reduced overall T-cell proliferation and release (**Fig. S2C**). Given our goal of designing a hydrogel depot that supports sustained T-cell expansion and release, we proceeded with unmodified NPs in subsequent studies.

To determine whether cytokines influence T-cell phenotype, we collected ACTIVATE hydrogels both three days and eight days post-encapsulation, dissociated them, and performed flow cytometric analysis. Although IL-12 did not enhance proliferation, it significantly preserves antigen sensitivity, as indicated by elevated PD-1 expression, up to eight days post-encapsulation compared to the other cytokines (**Fig. 2H-I, Fig. S3D**). In terms of memory phenotype, all cytokines maintained the effector memory phenotype of the stimulated PMEL T cells after three days of encapsulation (**Fig. 2J**). Similar frequencies of effector memory phenotypes were also observed after eight days of encapsulation (**Fig. S3C**). Interestingly, IL-7 progressively promoted a central memory phenotype over time (**Fig. 2J, Fig. S3C**), consistent with its well-characterized role in expanding naïve and central memory T-cell pools^37^. Together, these results demonstrated that ACTIVATE serves as a robust and adaptable platform for cytokine-mediated modulation of T-cell proliferation and phenotype. While IL-2 and IL-7 strongly promoted T-cell expansion, IL-12 uniquely preserved antigen sensitivity, highlighting the potential of this hydrogel system for tailoring T-cell responses in immunotherapy applications.

### 2.2. IL-12 ACTIVATE significantly delays B16-F10 tumor growth by engaging the endogenous immune system

Given that each interleukin uniquely influenced T-cell proliferation and phenotype *in vitro*, we next sought to determine whether these differences translated into distinct therapeutic efficacies *in vivo*. To assess the capacity of ACTIVATE to engage both transferred T cells and the endogenous immune system, we employed both the B16-F10 melanoma model and the E.G7-Ova lymphoma models in C57Bl/6 mice. Following *in vitro* stimulation and activation, PMEL CD8^+^ T cells were administered either intravenously (IV) or via peritumoral (PT) injection of ACTIVATE in animals induced with B16-F10 cells to generate an aggressive, poorly immunogenic melanoma model^38^ (**Fig. 3B-D**). Similarly, stimulated and activated OT-1 CD8^+^ T cells were administered either IV or via PT ACTIVATE in E.G7-Ova tumor-bearing mice **(Fig. 3E-G**). Mice were inoculated subcutaneously with either 3×10^5^ B16-F10 cells or E.G7-Ova cells and treated seven days post-inoculation with a 100μL PT dose of ACTIVATE containing 5×10^6^ adoptive PMEL cells (for B16-F10) or OT-1 T cells (for E.G7-Ova) and 2.5μg of the respective interleukin (**Fig. 3A**). Treatment-induced toxicity was monitored by assessing weight loss over one week post-treatment. The IV control group received systemically administered PMEL or OT-1 T cells with an equivalent dose of IL-2, which is commonly used to support the survival and expansion of adoptively transferred PMEL in preclinical models^39–41^.

**Figure 3.**
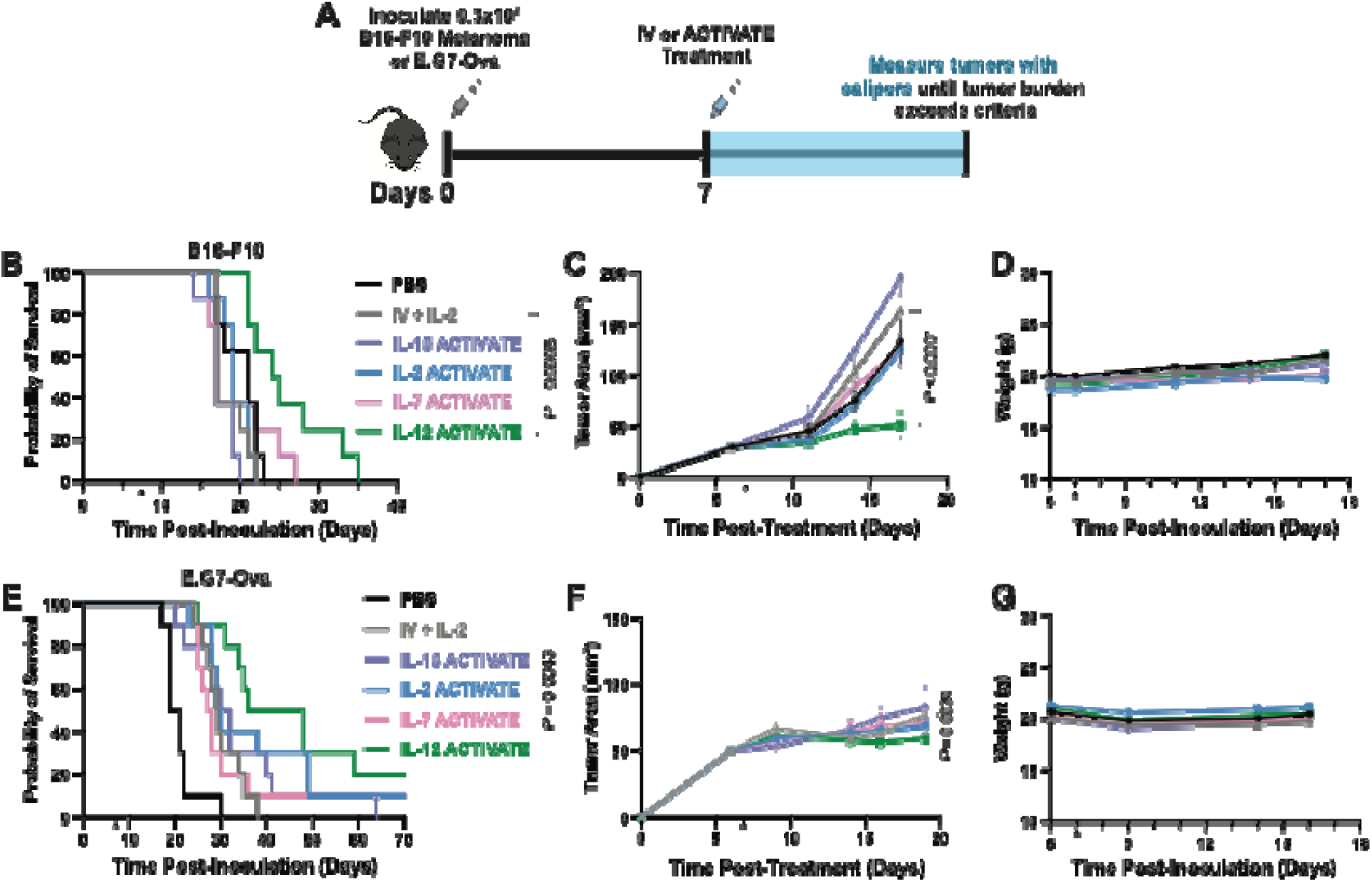
IL-12 ACTIVATE significantly delays B16-F10 and E.G7-Ova tumor growth. (**A**) C57Bl/6 mice were inoculated (s.c.) with 0.3×10^6^ B16-F10 cells (B-D) or 0.3×10^6^ E.G7-Ova (E-G) on D0 and treated on D7 with 100μL of ACTIVATE (p.t.) containing 5×10^6^ PMEL T cells (B-D) or 5×10^6^ OT-1 T cells (E-G) and 2.5μg of either IL-2, IL-7, IL-15, or IL-12. I.V. control group received 5×10^6^ PMEL or OT-1 T cells and 2.5μg of IL-2 administered retro-orbitally in a 100μL bolus. Tumors were measured with digital calipers until tumor burden exceeded euthanasia criteria. (**B**) Overall survival of treated and untreated mice for B16-F10. (**C**) Primary tumor growth curve for B16-F10. (**D**) Mouse weights after ACTIVATE or IV treatment (indicated by arrows) in B16-F10 study. (**E**) Overall survival of treated and untreated mice for E.G7-Ova. (**F**) Primary tumor growth curve for E.G7-Ova. (**G**) Mouse weights after ACTIVATE or IV treatment (indicated by arrows) in E.G7-Ova study. Data are representative of *n* = 8 (B-D) or *n* = 10 (E-G) and presented as mean ± s.e.m. *P* values were determined using the log-rank (Mantel-Cox) test (**C, F**) or two-way ANOVA followed by Tukey’s multiple comparison test (**B, E**) using GraphPad PRISM.

Interestingly, PT administration of IL-12 ACTIVATE significantly enhanced tumor control compared to all other treatments in both ACT models, raising the median survival time from 17 days for the IV control group to 24.5 days in the B16-F10 tumor-bearing mice (***P* = 0.0008, Fig. 3B-C**), and from 29 days for the IV group to 42 days in the E.G7-Ova tumor-bearing mice (***P* = 0.0043, Fig. 3E-F, Supplementary Table 1**). This outcome is consistent with the broad immunostimulatory effects elicited by IL-12, which extend beyond direct T-cell expansion, including stimulation of both innate and adaptive immune mechanisms^31^. Notably, none of the treatment groups exhibited significant weight loss within one week post-treatment, indicating a favorable safety profile (**Fig. 3D&G**). Thus, while IL-12 exhibited minimal impact on T-cell proliferation *in vitro*, its incorporation into ACTIVATE dramatically improved tumor control *in vivo*, underscoring the capacity of cytokine-loaded hydrogel depots to shape immune responses beyond direct effects on adoptively transferred cells.

ACT has been significantly enhanced by lymphodepletion, a standard clinical procedure involving sub-lethal irradiation that improves T-cell engraftment and peristence^42^. Given the efficacy of IL-12 ACTIVATE in the absence of lymphodepletion, we sought to determine whether preconditioning could further enhance the engraftment of PMEL cells delivered via ACTIVATE. To test this, we repeated the survival study with the key modification that mice underwent lymphodepletion one day prior to treatment. Interestingly, while lymphodepletion uniformly shifted the median survival of all treatment groups—regardless of IV or ACTIVATE administration—by approximately two days, it only significantly improved survival in the IL-15 ACTIVATE group (***P* = 0.004, Fig. S4, Supplementary Table 2**). In stark contrast, lymphodepletion abrogated the therapeutic efficacy of IL-12 ACTIVATE, as evidenced by markedly diminished tumor control (**Fig. S4B**). Clinically, lymphodepletion is associated with substantial toxicity and can likely exacerbate adverse effects of cytokine-based therapies^43^ . Consistent with these observations, all treated mice experienced >5% weight loss, and one mouse succumbed to treatment-related toxicity (**Fig. S4A, S4E**), likely due to the depletion of endogenous cytokine sinks potentiating the systemic effects of delivered interleukins^44^. These findings underscore that lymphodepletion is not required to enhance T-cell engraftment with ACTIVATE, and in fact, an intact immune system can synergize with the hydrogel-based therapy to achieve superior anti-tumor efficacy. Accordingly, all subsequent studies are conducted without lymphodepletion preconditioning unless otherwise specified.

### 2.3. ACTIVATE boosts adoptive and endogenous T-cell cytotoxicity and recruitment to tdLNs

Given the robust *in vivo* efficacy of IL-12 ACTIVATE, we sought to investigate the impact of ACTIVATE on both adoptively transferred cells and the host immune system. Specifically, we focused on characterizing endogenous immune responses elicited by IL-12 ACTIVATE, given its superior anti-tumor efficacy, and IL-2 ACTIVATE, due to its role in promoting T-cell proliferation *in vitro* and its frequent administration alongside PMEL in other studies^39^. The effects of these two treatments were compared to systemic PMEL and IL-2 administration.

To assess these effects, C57Bl/6 mice were subcutaneously inoculated with 3×10^5^ B16-F10 tumor cells and treated seven days later with either 100μL of PT administered ACTIVATE or systemic IV administration. Seven days post-treatment, tdLNs were harvested and analyzed via flow cytometry (**Fig. 4A**). Interestingly, ACTIVATE treated groups exhibited significantly higher numbers of cells in the tdLNs compared to the IV control (**Fig. 4B**). Concurrently, both IL-2 ACTIVATE and IL-12 ACTIVATE increased the expansion of PMEL cells and recruitment of endogenous immune cells found in the tdLNs (**Fig. 4C**).

**Figure 4.**
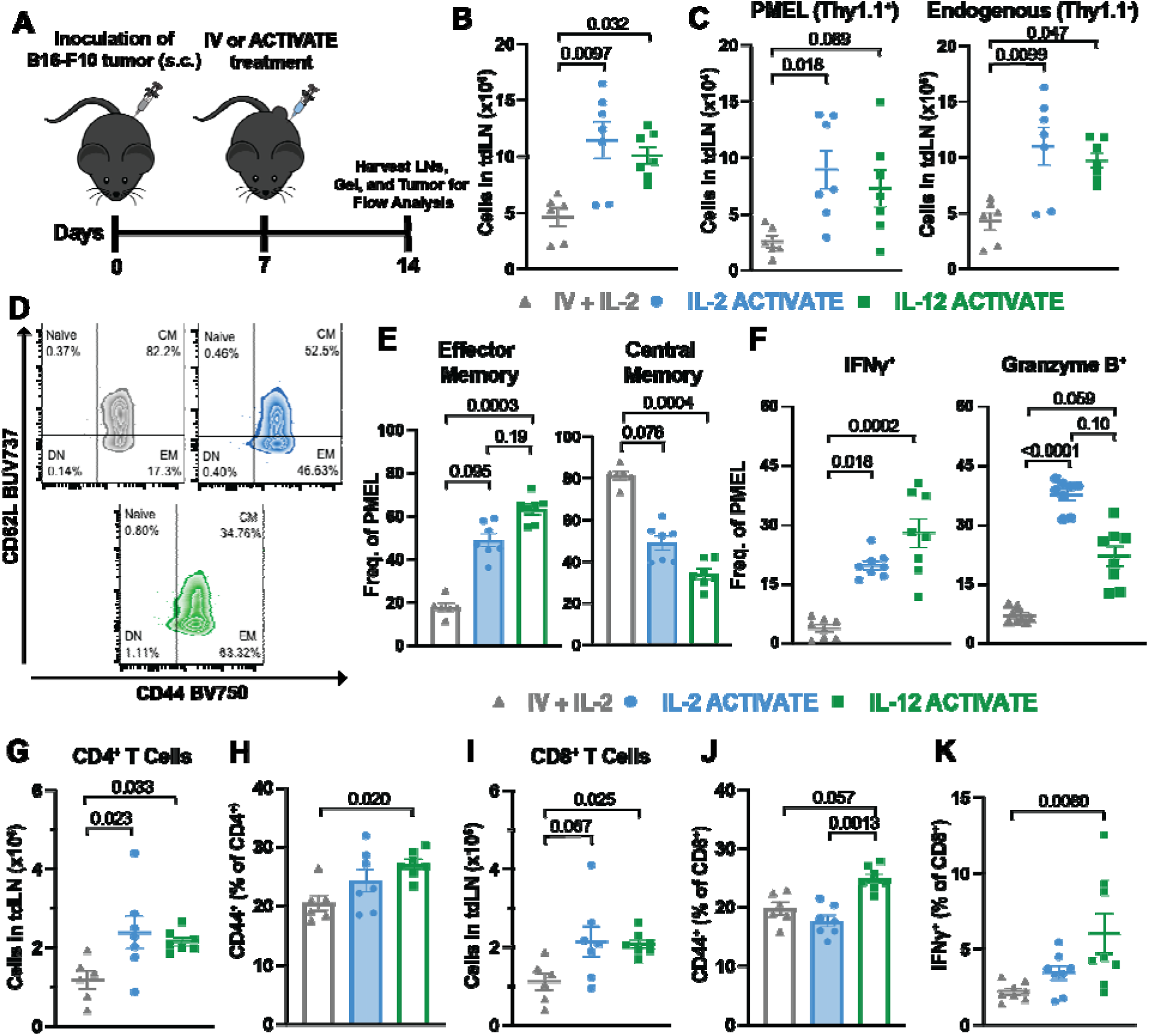
ACTIVATE promotes T-cell recruitment and cytotoxicity to the tdLNs. (**A**) Timeline schematic of experimental design: C57Bl/6 mice were inoculated (s.c.) with 0.3×10^6^ B16-F10 cells on D0 and treated on D7 with 100μL of ACTIVATE (p.t.) containing 5×10^6^ PMEL T cells and 2.5μg of either IL-2 or IL-12. 7 days after treatment (D14), tumor-draining lymph nodes were harvested and stained for flow cytometry analysis. (**B**) Number of total live cells (**C**) and total PMEL (Thy1.1^+^CD3^+^) and endogenous (Thy1.1^-^) cells found in the tdLNs for each respective treatment group. (**D**) Representative day 7 flow plots and (**E**) quantification of effector memory (CD62L^-^CD44^+^) and central memory (CD62L^+^CD44^+^) frequencies among PMEL cells in the tdLNs. (**F**) Frequency of IFNγ and Granzyme B populations among PMEL cells. (**G**) Total CD4^+^ T cells found in tdLNs of respective treatment groups and (**H**) frequency of CD44^+^ populations among CD4^+^ cells. (**I**) Total CD8^+^ T cells in tdLNs and (**J**) frequency of CD44^+^ populations and (**K**) IFNγ^+^ populations among CD8^+^ T cells. Data are representative of *n* = 6 to 8 mice per condition from two independent experiments and presented as mean ± s.e.m. *P* values were determined using Kruskal-Wallis test with Dunn’s multiple comparisons test using GraphPad PRISM.

Given the distinct effects of co-encapsulated interleukins on the adoptive T-cell phenotype *in vitro*, we wondered whether similar maintenance of memory phenotype would be observed *in vivo*. We were particularly interested in whether the adoptive cells that homed to the lymph nodes maintained their effector memory phenotype. Indeed, both ACTIVATE groups resulted in a higher frequency of effector memory PMEL cells, whereas the IV group gives rise to a greater frequency of central memory PMEL cells (**Fig. 4D-E**). IL-12 based IFNγ signaling has been previously found to be critical to adoptive cell-mediated anti-tumor activity^45^. Congruently, both ACTIVATE treatments enhanced the cytotoxic profile of the adoptive cells, with IL-12 ACTIVATE inducing a higher frequency of IFNγ-expressing PMEL cells (**Fig. 4F**). Further, both gels significantly increased the proportion of Granzyme B-expressing PMEL cells compared to systemically administered cells (**Fig. 4F**). These findings suggest that ACTIVATE not only enhances adoptive T-cell migration to the tdLNs but also bolsters their effector functionality.

Since interleukins can modulate endogenous T-cell responses, we further investigated whether ACTIVATE influences endogenous T-cell recruitment to the tdLNs. Notably, both IL-12 and IL-2 ACTIVATE significantly increased the numbers of CD4^+^ and CD8^+^ T cells in the tdLNs (**Fig. 4G & 4I**). However, only IL-12 ACTIVATE induced a marked elevation in CD44 expression on both CD4^+^ and CD8^+^ T cells relative to the systemic control and IL-2 ACTIVATE group (**Fig. 4H & 4J**), suggesting heightened activation. Furthermore, IL-12 ACTIVATE further enhanced CD8^+^ T-cell cytotoxicity, as evidenced by elevated IFNγ expression (**Fig. 4K**). Consistent with observations of the tdLN compartment, analysis of splenic T cells at the same time point revealed increased CD44 expression on both CD4^+^ and CD8^+^ T cells of IL-12 ACTIVATE treated mice, indicating systemic immune activation (**Fig. S6D**). These results demonstrate that ACTIVATE, particularly IL-12 ACTIVATE, promotes robust expansion and enhance effector functionality of adoptive T cells, while also stimulating endogenous T-cell recruitment and activation in the tdLNs, collectively contributing to a more potent anti-tumor immune response.

### 2.4. ACTIVATE enhances activation and recruitment of APCs to the gels and tdLNs

We have previously demonstrated that the dynamic nature of the hydrogel material enables simultaneous recruitment of immune cells into the PNP hydrogel in the context of vaccines, creating a local inflammatory niche that enhances the magnitude and duration of immune responses^27,46–48^. We hypothesized that a similar phenomenon may be occurring with the ACTIVATE hydrogels, facilitating immune cell recruitment and amplifying antitumor responses. To test this, we excised the ACTIVATE hydrogels seven days post-treatment, dissociated them, and analyzed their cellular infiltrates via flow cytometry (**Fig. 4A**, **Fig. 5A**).

**Figure 5.**
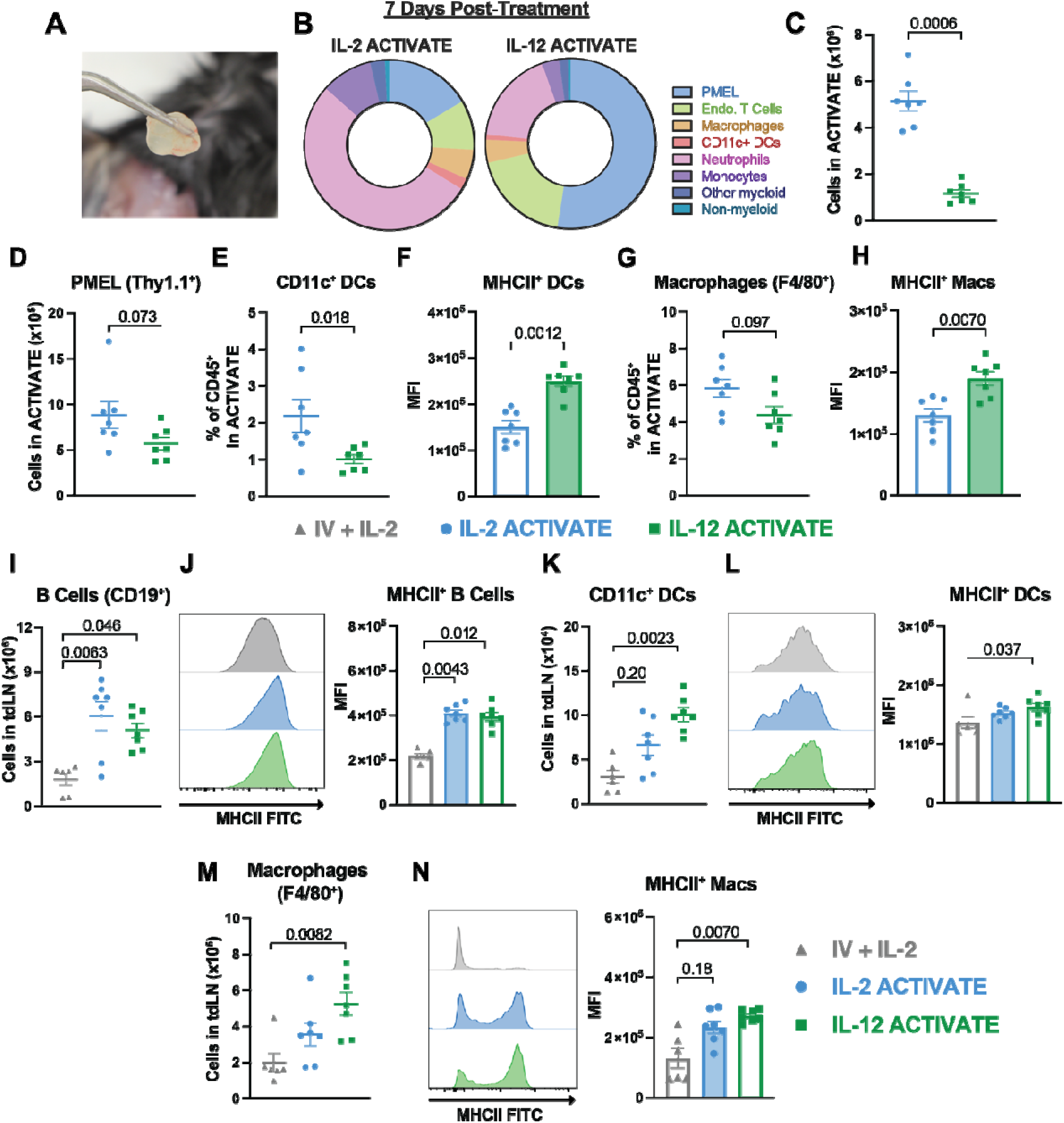
ACTIVATE enhances activation and recruitment of antigen-presenting cells to both the gels and the tdLNs. C57Bl/6 mice were inoculated (s.c.) with 0.3×10^6^ B16-F10 cells on D0 and treated on D7 with 100μL of ACTIVATE (p.t.) containing 5×10^6^ PMEL T cells and 2.5μg of either IL-2 or IL-12. One week after treatment (D14), tumor-draining lymph node (tdLN) and hydrogels were harvested and stained for flow cytometry analysis. (**A**) Picture of surgically removed ACTIVATE hydrogel 7 days post-treatment *in vivo*. (**B**) The frequency of PMEL adoptive T cells, endogenous T cells, macrophages, dendritic cells, neutrophils, monocytes, other myeloid cells, and non-myeloid cells within the CD45^+^ cell populations found in the IL-2 and IL-12 hydrogels. (**C**) Total number of live cells and (**D**) PMEL cells found in each respective hydrogel. (**E**) Frequency of CD11c^+^ DCs and (**G**) macrophages (F4/80^+^) found in the ACTIVATE hydrogels. (**F, H**) MFI quantification of MHCII^+^ expression on CD11c^+^ DCs (**F**) and macrophages (**H**) found in each respective hydrogel. Data are representative of *n* = 7 mice per condition and presented as mean ± s.e.m. *P* values were determined using Mann-Whitney test using GraphPad PRISM. (**I**) Total number of B cells (CD19^+^) found in tdLNs. (**J**) Representative day 7 flow plots and quantification of MHCII^+^ B cells. (**K**) Total number of CD11c^+^ DCs found in tdLNs. (**L**) Representative day 7 flow plots and quantification of MHCII^+^ CD11c^+^ DCs. (M) Total number of macrophages (F4/80^+^) found in tdLNs. (**N**) Representative day 7 flow plots and quantification of MHCII^+^ macrophages. Data are representative of *n =* 6 for IV PMEL + IL-2 treatment group and *n =* 7 for respective gel groups and presented as mean ± s.e.m. *P* values were determined using Kruskal-Wallis test with Dunn’s multiple comparisons test using GraphPad PRISM.

Remarkably, both ACTIVATE hydrogels recruited a diverse immune cell repertoire, including endogenous T cells, macrophages, neutrophils, dendritic cells, and more (**Fig. 5B**). While IL-12 ACTIVATE recruited a similar number of cells to what we previously observed in vaccine settings^27^, IL-2 ACTIVATE exhibited a ∼3-fold increase in total cellular recruitment (**Fig. 5C**). We observed IL-2 ACTIVATE niches contained significantly higher infiltration of endogenous immune cells, natural killer (NK) cells, neutrophils, monocytes, and other myeloid populations, while also retaining a greater number of adoptive cells at this same time point (**Fig. S7B, Fig. 5D**). Most notably, although IL-2 ACTIVATE recruited higher frequencies and absolute counts of CD11c^+^ dendritic cells (DCs) and F4/80^+^ macrophages, IL-12 ACTIVATE distinctively upregulated MHC-II expression on both APC subsets, indicating enhanced antigen presentation and heightened immune activation (**Fig. 5E-F, 5H**).

Given the robust immune activation within the transient ACTIVATE hydrogel niche, we next examined whether ACTIVATE influences APC recruitment and activation in the tdLNs. Both IL-2 and IL-12 ACTIVATE significantly increased CD19^+^ B cell recruitment to the tdLNs while also enhancing their MHC-II expression relative to systemically administered controls (**Fig. 5I-J**). A similar trend was seen in splenic APCs, where both hydrogels expanded the population of antigen-presenting B cells (**Fig. S6A**). While both ACTIVATE increased CD11c^+^ DC recruitment to the tdLNs, only IL-12 ACTIVATE significantly upregulated MHCII expression on these DCs (**Fig. 5K-L**). Further analysis of splenic DCs revealed that IL-12 ACTIVATE preferentially expanded cDC1s, which prime CD8^+^ T cell responses and drive type-1 immunity, whereas systemic adoptive cell and cytokine administration skewed DC populations towards cDC2s, which preferentially activate CD4^+^ T cells and promote Th2/Th17 responses^49^ (**Fig. S6B-C**). Lastly, IL-12 ACTIVATE not only recruited significantly more macrophages to the tdLNs but also enhanced their activation status, as reflected by elevated MHC-II expression (**Fig. 5M-N**). Thus, ACTIVATE established a robust inflammatory niche that recruits diverse immune cell subsets, with IL-2 ACTIVATE preferentially driving overall immune infiltration while IL-12 ACTIVATE enhanced antigen presentation. These findings underscore the ability of ACTIVATE to modulate both local and systemic immune landscapes, ultimately shaping distinct antitumor immune responses.

We previously demonstrated that co-encapsulation of IL-15 within the PNP hydrogel drives CAR-T cell expansion, allowing the material to function as a transient inflammatory niche for sustained T-cell proliferation beyond the initially loaded dose^25^. Since these studies were conducted in immunocompromised mice, we sought to understand the influence of immune pre-conditioning on recruitment of cells to ACTIVATE. To directly compare immune recruitment dynamics, we subcutaneously inoculated mice with B16-F10 melanoma cells and treated them after seven days with PT administration of IL-2 ACTIVATE. To assess the role of endogenous immune cells, one treatment cohort underwent lymphodepletion one day prior to treatment. Gels were excised three days post-treatment, dissociated, and analyzed for infiltrate composition.

As predicted, ACTIVATE hydrogels from lymphodepleted mice exhibited significantly higher numbers of adoptive PMEL cells, whereas non-lymphodepleted gels showed increased recruitment of endogenous cells (**Fig. S8A-B**). While frequencies and counts of endogenous T cells, neutrophils, and NK cells were similar between conditions, gels from non-lymphodepleted mice contained significantly greater numbers of CD11c^+^ DCs, other myeloid cells, and non-myeloid cells (**Fig. S8C**). These results indicate that ACTIVATE dynamically adapts to the host immune status, with lymphodepletion favoring adoptive T-cell expansion while an intact immune system supports a broader and synergistic endogenous response. These observations underscore the versatility of ACTIVATE as a platform capable of shaping immune responses based on the surrounding immune landscape.

### 2.5. ACTIVATE enhances tumor infiltration and cytotoxicity of effector immune cells

To evaluate how ACTIVATE reshapes the TME, we harvested, dissociated, and analyzed tumors via flow cytometry seven days post-treatment as previously described (**Fig. 4A**). We discovered that both ACTIVATE formulations recruited significantly more endogenous immune cells to the tumor compared to systemic PMEL administration (**Fig. 6A**). While the total number of adoptive PMEL cells in the tumor remained similar across all groups at this timepoint, ACTIVATE—particularly the IL-12 formulation—enhanced the cytotoxic function of intratumoral PMEL cells. A significantly higher frequency of IFN ^+^GzmB^+^ PMEL cells was observed for the IL-12 ACTIVATE treatment, mirroring the heightened activation observed in the tdLNs, as well as increased IFNγ expression compared to the IV group (**Fig. 6B-D**).

**Figure 6.**
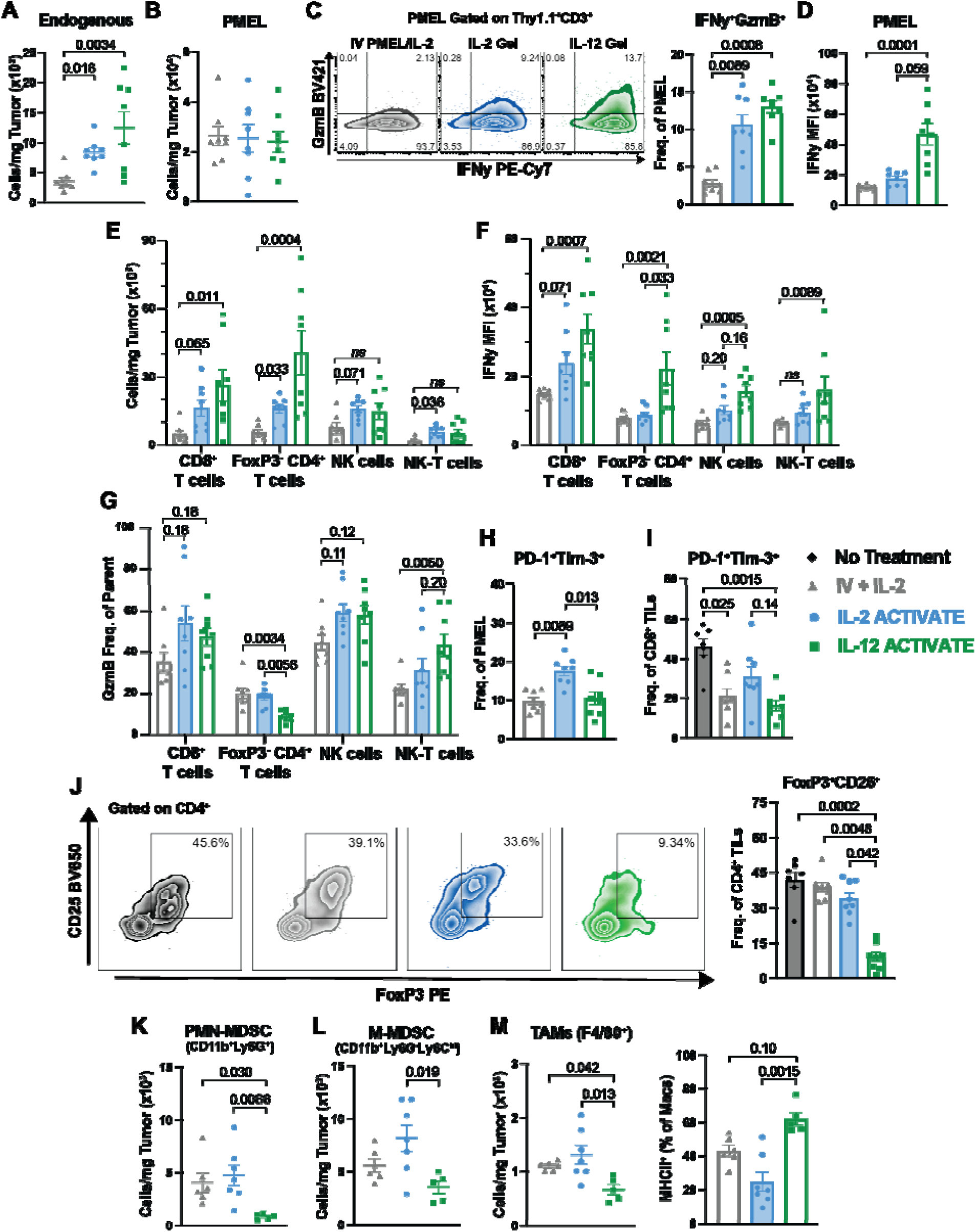
ACTIVATE enhances recruitment of effector lymphocytes to tumors and boosts their cytotoxicity. C57Bl/6 mice were inoculated (s.c.) with 0.3×10^6^ B16-F10 cells on D0 and treated on D7 with 100μL ACTIVATE (p.t.) containing 5×10^6^ PMEL T cells and 2.5μg of either IL-2 or IL-12. 7 days after treatment (D14), tumors were harvested and stained for flow cytometry analysis. (**A**) Counts of endogenous (Thy1.1^-^) cells and (**B**) PMEL (Thy1.1^+^CD3^+^) cells found in tumors harvested 7 days after treatment. (**C**) Representative flow plot and frequencies of GranzymeB^+^IFN ^+^ amongst PMEL cells. (D) MFI of IFN ^+^ PMEL cells in the tumor. (**E**) Counts of tumor-infiltrating CD8^+^ T cells, FoxP3^-^CD4^+^ T cells, NK cells (NK1.1^+^), and NK-T cells (NK1.1^+^CD3^+^) found in the tumor on D14. (**F**) MFI of IFNγ expressing CD8 T cells, CD4^+^ T cells, NK cells, and NK-T TILs. (**G**) Frequency of granzyme B^+^ expressing CD8^+^ T cells, CD4^+^ T cells, NK cells, and NK-T TILs. **(H, I)** Frequency of terminally exhausted (PD-1^+^Tim-3^+^) PMEL (**H**) and endogenous CD8^+^ (**I**) tumor-infiltrating lymphocytes. **(J)** Representative flow cytometry plots and frequencies of Regulatory T cells (FoxP3^+^CD25^+^) gated on CD4^+^ T cells. (**K)** Counts of PMN-MDSC (CD11b^+^Ly6G^+^) and (**L**) M-MDSC (CD11b^+^Ly6G^-^Ly6C^hi^) found in the tumor on D7. (**M**) Counts of tumor-associated macrophages (F4/80^+^) in the tumor on D7 and frequency of MHCII^+^ expressing macrophages for each respective treatment group. Data are representative of *n* = 8 for all treatment groups and presented as mean ± s.e.m. *P* values were determined using Kruskal-Wallis test with Dunn’s multiple comparisons test using GraphPad PRISM.

In addition to boosting PMEL cell function, both ACTIVATE formulations significantly increased the recruitment of diverse tumor-infiltrating leukocytes (TILs) compared to IV administration. These observations included a marked rise in CD8^+^ cytotoxic T cells and FoxP3^-^CD4^+^ helper T cells in the TME (**Fig. 6E**). Interestingly, IL-2 ACTIVATE also enriched NK cells and NK-T cells in the tumor, consistent with the well-documented role of IL-2 in supporting their proliferation^50^ (**Fig. 6E**). Importantly, both ACTIVATE formulations enhanced IFNγ expression and Granzyme B production across multiple TIL subsets, indicating a broad increase in cytotoxic effector function within the TME (**Fig. 6F-G**). Collectively, these findings demonstrate that ACTIVATE gels not only potentiate the function of adoptively transferred T cells but also drive recruitment and activation of diverse endogenous immune effectors, reinforcing a highly inflammatory TME.

### 2.6. IL-12 ACTIVATE reduces immunosuppressive mechanisms in the TME

Given the potent antitumor effects observed with IL-12 and its known pleiotropic role in reducing regulatory T cells (T_REGS_) and mitigating T-cell exhaustion^51,52^, we next assessed its impact on immunosuppressive populations and T-cell exhaustion within the TME. Flow cytometric analysis revealed that IL-12 ACTIVATE reduced the frequency of terminally exhausted PD-1^+^Tim-3^+^ PMEL cells and endogenous CD8^+^ TILs compared to IL-2 ACTIVATE and untreated controls (**Fig. 6H-I**). This reduction in terminally exhausted T cells suggests that IL-12 ACTIVATE has the potential to improve the persistence and antitumor function of transferred cells, ultimately enhancing the therapeutic impact of ACT. Although the standard IV treatment similarly decreased the frequency of terminally exhausted PMEL compared to IL-2 ACTIVATE and endogenous CD8^+^ TILs, these changes conferred a less pronounced therapeutic benefit compared to IL-12 ACTIVATE in vivo (**Fig. 3B-C**).

Remarkably, IL-12 ACTIVATE also leads to a significant reduction in intratumoral CD4^+^CD25^+^FoxP3^+^ T_REGS_, a highly immunosuppressive subset linked to poor prognosis and resistance to immunotherapy (**Fig. 6J**). Additionally, IL-12 ACTIVATE diminished absolute counts of polymorphonuclear myeloid-derived suppressor cells (PMN-MDSCs) compared to both IV and IL-2 ACTIVATE treatments, while also significantly reducing monocytic MDSCs (M-MDSCs) relative to IL-2 ACTIVATE (**Fig. 6K-L**). MDSCs have been implicated in promoting T cell dysfunction, dampening antitumor immunity, and supporting tumor progression by secreting immunosuppressive cytokines and reactive oxygen species^53^. The reduction observed here suggests that IL-12 ACTIVATE may counteract multiple suppressive mechanisms within the TME, thereby fostering a more immunostimulatory environment.

Finally, IL-12 ACTIVATE significantly decreased the presence of tumor-associated macrophages (TAMs) within the TME (**Fig. 6M**). Notably, the TAMs that remained exhibit an enhanced activation phenotype, with upregulated MHCII expression compared to both IL-2 ACTIVATE and IV-treated groups, suggesting a shift toward a more pro-inflammatory, antigen-presenting phenotype. Together, these findings highlight the dual impact of ACTIVATE gels: enhancing cytotoxic immune infiltration while simultaneously diminishing key immunosuppressive mechanisms within the TME. While IL-2 ACTIVATE was found to broadly recruit immune effectors, IL-12 ACTIVATE uniquely reshaped the TME by reducing suppressive cell populations, promoting antigen presentation, and amplifying effector cell cytotoxicity. These data highlight the versatility of ACTIVATE hydrogels as a precision immunotherapy platform capable of modulating both pro- and anti-tumor immune networks to drive robust antitumor responses.

### 2.7. Checkpoint blockade and cytokine combinations potentiate ACTIVATE-mediated tumor clearance

Given the exhaustion phenotype observed in both adoptive PMEL cells and endogenous CD8^+^ TILs, as well as the robust immune activation induced by ACTIVATE, we sought to determine whether this platform could synergize with immune checkpoint blockade, the current standard-of-care immunotherapeutic treatment for solid tumors^54^. IL-12 has previously demonstrated potent synergy with ICB^15,55^, making IL-12 ACTIVATE a compelling candidate for combination immunotherapy.

To test this, we repeated our prior *in vivo* studies, incorporating three doses of 250μg of anti-PD-1 administered intraperitoneally once every three days, beginning on the day of ACTIVATE or IV treatment (**Fig. 7A**). Tumor growth and survival analyses revealed that IL-12 ACTIVATE in combination with ICB significantly enhanced tumor clearance, with 50% of treated mice (5/10) remaining tumor-free by day 50 (**Fig. 7B, Supplementary Table 3**). To assess the durability of this response, we rechallenged these five mice 110 days after initial tumor inoculation by injecting 1×10^5^ B16-F10 melanoma cells on the contralateral flank. Remarkably, all five mice remained tumor-free by day 50, whereas age-matched, naïve control mice succumbed to tumor burden within 22 days (**Fig. 7C**). These findings indicate that IL-12 ACTIVATE, in combination with ICB, elicits durable anti-tumor immunity and establishes robust long-term immune memory.

**Figure 7.**
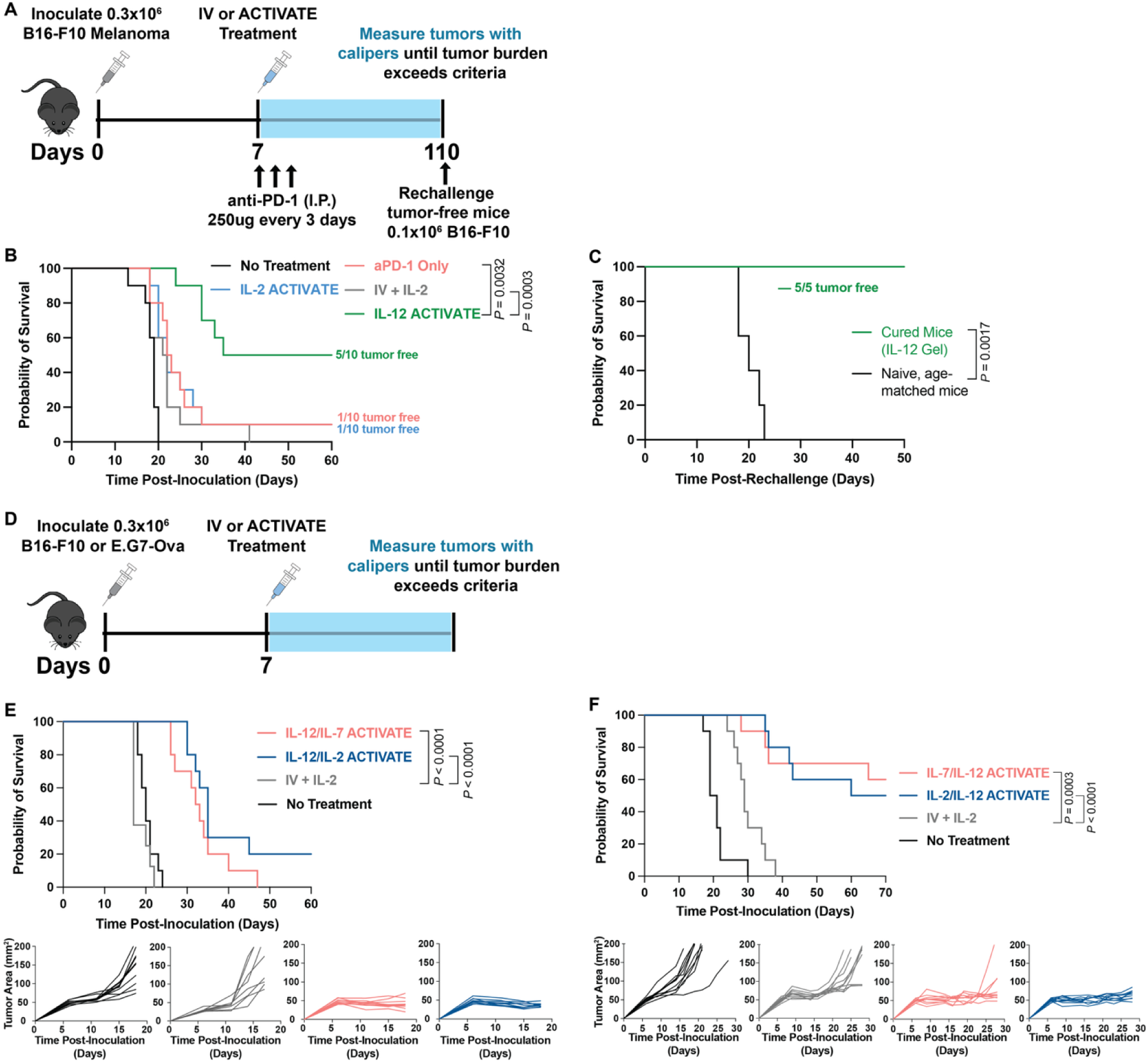
Checkpoint blockade and cytokine combinations potentiate ACTIVATE efficacy. (**A**) Experimental timeline: C57Bl/6 mice were inoculated (s.c.) with 0.3×10^6^ B16-F10 cells on D0 and treated on D7 with 100μL ACTIVATE (p.t.) containing 5×10^6^ PMEL T cells and 2.5μg of either IL-2 or IL-12. All treated mice received 250μg of anti-PD-1 checkpoint intraperitoneally once every 3 days beginning on day 7. Any mice that were cancer free were rechallenged on Day 110 with 0.1×10^6^ B16-F10 cells on the opposite flank. (**B**) Overall survival of the treated mice, and (**C**) overall survival of rechallenged mice. Treatments groups included no treatment, which received s.c. PBS, IV PMEL/IL-2, IL-2 gel, IL-12 gel, and anti-PD-1 only as an added control for (**B**) and cured mice from the IL-12 gel group as well as naïve, age-matched mice for (**C**). (**D**) Experimental timeline: C57Bl/6 mice were inoculated (s.c.) with either 0.3×10^6^ B16-F10 cells or E.G7-Ova cells on D0 and treated on D7 with 100μL ACTIVATE (p.t.) containing either 5×10^6^ PMEL T cells or OT-1 T cells, respectively, and 2.5μg of IL-12 and 2.5μg of either IL-2 or IL-7. I.V. control group received 5×10^6^ PMEL or OT-1 T cells and 2.5μg of IL-2 administered retro-orbitally in a bolus. Tumors were measured with digital calipers until tumor burden exceeded euthanasia criteria. (**E**) Overall survival of the treated mice and tumor growth curves for B16-F10 tumors. (**F**) Overall survival of the treated mice and tumor growth curves for E.G7-Ova tumors. Data are representative of *n* = 8 or 10 from two independent experiments and presented as mean ± s.e.m*. P* values were determined using the log-rank (Mantel-Cox) test (**C, E, F**) using GraphPad PRISM.

Given the modularity of ACTIVATE, which allows for facile incorporation of cytokines via simple mixing without requiring additional modification of cargo, we explored whether cytokine combinations could further enhance the anti-tumor efficacy of the treatment. Specifically, we investigated whether the proliferative benefits of IL-2 or IL-7 could be synergistically combined with the immune-activating properties of IL-12. To test this, we conducted survival studies against subcutaneous B16-F10 or E.G7-Ova similar to our previous experiments, incorporating 2.5μg of IL-12 with either 2.5μg of IL-2 or 2.5μg of IL-7 into each 100μL ACTIVATE dose with the appropriate adoptive cell type (**Fig. 7D**). In the B16-F10 model, both cytokine combinations significantly improved therapeutic efficacy, with the IL-12/IL-7 combination extending median survival to 32.5 days and the IL-12/IL-2 combination further extending median survival to 35 days (***P* < 0.0001, Fig. 7E, Supplementary Table 4**). Further, 2 of the 10 mice treated with IL-12/IL-2 ACTIVATE remained tumor-free by 60 days post-inoculation (**Fig. 7E**). Similarly in the E.G7-Ova model, both cytokine combinations also significantly improved therapeutic efficacy, with the IL-12/IL-7 combination extending median survival beyond 70 days, with 6 of the 10 mice remaining tumor-free by 70 days post-inoculation (***P* = 0.003**) while the IL-12/IL-2 combination further extending median survival to 65 days, with 5 of the 10 mice remaining tumor-free by 70 days post-inoculation (***P* < 0.0001, Fig. 7F, Supplementary Table 4**). These findings suggest that while proliferative cues alone are insufficient to drive durable tumor control, combining them with potent immune-activating signals can orchestrate a more effective anti-tumor response. This highlights the versatility of the ACTIVATE platform, which can be readily adapted to incorporate distinct cytokine formulations tailored to specific tumor contexts.

## 3. Discussion

Adoptive cell therapies have revolutionized the treatment of hematological cancers but remain hindered in treating solid tumors by inefficient T-cell trafficking, on-target off-tumor toxicities, and cytokine release syndrome. These challenges are exacerbated by systemic administration, which limits T-cell infiltration and expansion within the immunosuppressive tumor microenvironment. To overcome these barriers, we developed ACTIVATE, an injectable dynamic hydrogel depot technology that establishes a transient inflammatory niche for the localized co-delivery of adoptive T cells and stimulatory cytokines. This approach facilitates sustained adoptive cell retention, enhances immune infiltration, and modulates the TME to foster robust and durable anti-tumor immunity.

The unique architecture of ACTIVATE prolongs cytokine retention within the niche while permitting active motility of entrapped cells, sustaining T-cell viability and tumor-reactive phenotypes. By enabling prolonged retention of proliferative cytokines such as IL-2 and IL-7, ACTIVATE promotes continued expansion of adoptive cells while mitigating toxicities associated with systemic cytokine administration. Moreover, our data indicates that proliferation alone is insufficient to drive consistent tumor clearance. Indeed, while IL-2 and IL-7 promoted robust T-cell expansion, their incorporation failed to yield durable tumor regression when delivered in isolation. In contrast, the addition of IL-12, a potent immune activator with minimal direct proliferative effect on T cells, significantly improved tumor control in the both the poorly immunogenic B16-F10 melanoma and E.G7-Ova solid tumor models—particularly when combined with IL-2 or IL-7—highlighting the importance of coordinating adoptive cell expansion with broader activation of the endogenous immune system.

Beyond augmenting adoptive T-cell function, ACTIVATE orchestrates systemic immune activation by recruiting and activating endogenous immune cells in the spleen, tumor-draining lymph nodes, tumors, and the hydrogel itself. The recruitment of professional antigen-presenting cells, including B cells, macrophages, and dendritic cells to the lymph nodes and the hydrogel amplify antigen presentation and enhance endogenous immune responses. IL-12 ACTIVATE further reprograms the TME by depleting immunosuppressive populations such as regulatory T cells, myeloid-derived suppressor cells, and tumor-associated macrophages, while simultaneously increasing the infiltration and cytotoxic activity of natural killer cells, CD4^+^ helper T cells, and CD8^+^ cytotoxic T cells. In this way, ACTIVATE mobilizes synergistic interaction between adoptive and endogenous immune cells in enhancing anti-tumor efficacy.

Cytokine dosing remains a critical determinant in optimizing immunotherapeutic efficacy, necessitating a precise balance between immune potentiation and toxicity mitigation. In this study, we utilized a 2.5μg cytokine dose extrapolated from clinical IL-2 regimens^39,56–59^ (**Supplementary Discussion**); however, our prior work demonstrated that lower concentrations, such as 0.25μg of IL-15, effectively promoted CAR-T cell expansion^25^. This observation suggests that cytokine dosing within ACTIVATE can be further refined, offering a tunable approach to achieving maximal immune modulation with minimized toxicity.

While this study utilized murine adoptive T cells as a model for TCR-based adoptive cell therapies, translating these findings to human T cells and CAR-T applications remains a crucial next step. Future work will require more immunologically competent settings, including humanized mouse models or murine CAR-T systems, to fully assess ACTIVATE’s ability to coordinate both adoptive and endogenous immune responses as well as its efficacy in long-term tumor control, particularly in abscopal and metastatic disease settings. Furthermore, validation of ACTIVATE in orthotopic and metastatic tumor models, exploration of its post-resection applications, and integration of immune-modulating antibodies to enhance adoptive T-cell efficacy will be required to more broadly evaluate the utility of ACTIVATE in various indications. Likewise, a more comprehensive interrogation of ACTIVATE’s ability to remodel systemic immune landscapes will be imperative for defining its role in generating long-term immunity against cancer. The flexibility of the ACTIVATE platform in delivering distinct cytokine formulations further broadens its applicability across diverse malignancies, enabling precision-engineered immune interventions tailored to tumor-specific pathophysiology.

Overall, ACTIVATE represents a next-generation, highly tunable cytokine and adoptive cell delivery system designed to circumvent critical limitations of ACT. By combining localized adoptive cell and cytokine delivery with sustained immune activation, it offers a transformative strategy for enhancing the efficacy and safety of ACT-based immunotherapies, paving the way for more durable and effective cancer treatments.

## 4. Materials and Methods

### Materials

All chemicals, reagents, and solvents used to make the PNP hydrogels were purchased as reagent grade from Sigma-Aldrich and used as received unless otherwise specified. Glassware and stir bars were oven-dried at 180°C.

### HPMC-C_12_ Synthesis

HPMC-C_12_ was prepared according to previously reported procedures^27^. Hydroxypropyl methylcellulose (HPMC) (1.0g) was dissolved in anhydrous N-methylpyrrolidone (NMP, 45mL) by stirring at 80°C for 1 hour. After dissolution, the solution was cooled to room temperature. A separate solution of 1-dodecyl isocyanate (105mg, 0.5mmol) in NMP (5mL) was prepared and added dropwise to the polymer solution, followed by the addition of N,N-diisopropylethylamine (Hunig’s base) (125uL) as a catalyst. The reaction mixture was stirred at room temperature for 16 hours before the modified polymer was precipitated in acetone. The resulting HPMC-C_12_ was collected via filtration, redissolved in Milli-Q water (approximately 2wt%), and purified by dialysis (MWCO 3kDa) against water for four days. The final product was lyophilized to yield a white amorphous solid and reconstituted in sterile 1x PBS (6wt%) for further use.

### Preparation of PEG-PLA and PEG-PLA NPs

PEG-*b*-PLA was synthesized following established protocols^27^. Prior to synthesis, diazabicyclo[5.4.0]undec-7-ene (DBU) was purified by distillation under reduced pressure and stored over molecular sieves until use, commercial lactide was recrystallized three times from ethyl acetate directly before use, PEG (methyl ether) was dried at 100°C under vacuum for 3 hours directly before user, and dichloromethane (DCM) was dispensed from a solvent purification system immediately before use. PEG (methyl ether) (2.5g of 5kDa) and DBU (15uL, 0.1mmol) were dissolved in anhydrous DCM (1mL) under an inert nitrogen atmosphere. Separately, lactide (1.0g, 6.9mmol) was also dissolved in DCM (3mL) under an inert nitrogen atmosphere. The lactide solution was rapidly added to the PEG/DBU solution and allowed to stir for 8 minutes. The reaction mixture was then quenched with aqueous acetic acid and precipitated by 1:1 hexane and ethyl ether solution. The final product was collected by centrifugation and dried under vacuum. Gel permeation chromatography (GPC) in DMF was used to verify the molecular weight and dispersity of polymers matched previously reported values.

NPs were prepared as previously reported^27^. PEG-PLA (50mg) was dissolved in 75:25 dimethyl sulfoxide (DMSO):acetonitrile (ACN) (1mL) and added dropwise to 10mL of water under high stir rate (600rpm) at room temperature. NPs were purified via centrifugation (4500 RCF) in centrifugal filter (Amicon Ultra-15, MWCO 10kDa) for 1 hour, then resuspended in 1x PBS to a final concentration of 20wt%. NPs were characterized by dynamic light scattering (DLS).

### PNP Hydrogel Formulation

Throughout this manuscript, a 1wt% HPMC-C12 and 5wt% PEG-PLA formulation was used for the ACTIVATE depots per previous optimization^25^. To prepare the PNP hydrogels, HPMC-C12 was dissolved at 6wt% in 1xPBS and loaded into a 1mL luer-lock syringe. A 20wt% solution of PEG-PLA NPs in 1x PBS was added to a solution of PBS with cytokine and/or adoptive cells, depending on formulation, and loaded into a second 1mL luer-lock syringe. These syringes were connected using a female-female luer-lock elbow and gently mixed until a homogenous PNP hydrogel was formed. Each ACTIVATE dose consisted of a 100μL injection peritumoral through a 23-guage needle. Rheological testing of the PNP hydrogel was performed (without cells) as previously described^25^.

### Mice, Cancer Cell Lines, and Cytokines

Seven- to eight-week-old female C57Bl/6J mice were purchased from The Jackson Laboratory. TCR-transgenic Thy1.1+ pmel-1 (PMEL) mice (B6.Cg-Thy1a/Cy Tg(TcraTcrb)8Rest/J) and OT-I mice (C57BL/6-Tg(TcraTcrb)1100Mjb/J) were also purchased from The Jackson Laboratory. Seven- to eight-week-old female Nod.Cg-*Prkdc*^scid^ *Il2rg*^tm1Wjl^/SzJ (NSG) mice were purchased from The Jackson Laboratory. All mice were housed in the animal facility at Stanford University (protocol – APLAC 33475). All the animal experiments performed in this research were approved by the Stanford Administrative Panel on Laboratory Animal Care. B16-F10 melanoma and E.G7-Ova cells were originally procured from the American Type Culture Collection and were cultured in RPMI medium (Gibco) containing 10% fetal bovine serum (FBS), penicillin (100 U/mL), and streptomycin (100μg/mL) and routinely tested for mycoplasma. Cells were cultured in a 5% CO_2_ environment at 37°C. Human cytokines were purchased from R&D Systems (Recombinant IL-2 Protein, Recombinant IL-7 Protein, Recombinant IL-2 Protein, Recombinant IL-15 Protein) and murine cytokines were purchased from Sino Biological (IL-2, IL-7, and IL-12A & IL-12B Heterodimer Protein (His Tag)) and PeproTech (Recombinant murine IL-15).

### Cytokine In Vitro Release

Glass capillary tubes (inner diameter = 3mm) were plugged at one end with epoxy, then incubated overnight at 5°C with a 0.1wt% solution of bovine serum albumin. After incubation, the tubes were dried using air and evaporation. 100μL of PNP hydrogel containing 0.5μg of either IL-2, IL-7, IL-12, or IL-15 was injected into the capillary tube, followed by the addition of 300μL of 1x PBS loaded on top of each gel. Samples were stored during assay at 37°C to simulate physiological conditions. At specified time points, the PBS was carefully aspirated using a long needle and stored at -80°C for later analysis before being replaced with fresh PBS. Cytokine concentrations were quantified using an enzyme-linked immunosorbent assay (ELISA) according to the manufacturer’s instructions (Human IL-2 DuoSet ELISA, Human IL-12 p70 DuoSet ELISA, Human IL-15 DuoSet ELISA, and Human IL-7 DuoSet ELISA; R&D Systems). Absorbance was measured at 450 nm using a Synergy H1 Microplate Reader (BioTek). At the end of the 7-day assay period, the gel was diluted 10-fold in saline and incubated for 3 hours before being stored for analysis of remaining cytokine concentration. Cytokine concentrations were determined based on standard curves. Mass of cytokine remaining in the gel was calculated as the difference between the total mass released during the study and the amount of cytokine remaining in the gel at the conclusion of the experiment.

### Preparation of PMEL and OT-1 Cells

Spleens and lymph nodes were harvested from either PMEL or OT-1 transgenic mice under sterile conditions. Single-cell suspensions were prepared by mechanically disrupting the spleen, followed by red blood cell lysis using ACK lysis buffer (Gibco). The isolated splenocytes were then plated at 100μL per well in a 96-well U-Bottom Plate (NEST) at a concentration of 2×10^6^ cells/mL in the presence of 60IU/mL of Recombinant Human IL-2 (PeproTech) and either 10μg/mL hGp100 (25-33) for PMEL or 10μg/mL Ova (257 – 264) Peptide Fragment (Anapsec Inc.) in complete RPMI 1640 (with glutamine, 1x non-essential amino acids, 1mM sodium pyruvate, 0.4x vitamin solution, 92μM 2-mercaptoethanol, 1% penicillin/streptomycin, and 10% FBS) and left to culture for 72 hours. If cells appeared overcrowded, cells were split and fed with half-media volume and 50IU/mL IL-2 and placed back in incubator. Cells were then harvested, centrifuged at 300 x g for 5 minutes, and plated at a concentration of 1×10^6^ cells/mL in fresh cRPMI supplemented with 60IU/mL IL-2 and 1μg/mL InVivoMab anti-CD28 (BioXCell) on 96-well plates incubated with 5μg/mL InVivoMab anti-mouse CD3 (BioXCell) and washed 3x with 1x PBS. Cells were incubated for 24 hours before being fed with half media volume and 120IU/mL IL-2. Cells were again incubated for another 48 hours before being harvested and moved to clean plates at a concentration of 1×10^6^ cells/mL in cRPMI supplemented with 60IU/mL IL-2. After ten days of stimulation and activation, cells were taken forward for in vitro and in vivo experiments.

### PMEL Cell Release Assay

PMEL T cells were incorporated into PNP hydrogel formulation at a concentration of 20 million cells/mL with 2.5μg of respective cytokine. A volume of 100μL of the hydrogel was then injected into a transwell insert (15mm Netwell Insert with 74μm Mesh Size) and carefully placed into a 12-well plate and 1.2mL of RPMI with no additives was gentle below and placed in a 5% CO_2_ incubator at 37°C. At each time point, the medium beneath the transwell was carefully removed, the cells in which were counted, and the transwell was placed into a new well with an equal volume of fresh media. At the end of the experiment (day 8), the remaining hydrogel contents of the transwell were diluted, the cells inside were counted, and cell viability was recorded. For in vitro phenotype studies, the cell release assay was prepared as described. Transwells were shifted to fresh media each day to prevent cell migration back into the hydrogel. Three days and eight days post-encapsulation, hydrogels were removed from transwells, mechanically dissociated and diluted, and cell count, and viability were recorded. These cells were then stained using flow cytometry methods described below.

### Murine Adoptive Cell Therapy Models

Seven- to ten-week old mice were shaved on their right flank and inoculated subcutaneously with 50μL of either B16-F10 or E.G7-Ova cells (6 x 10^6^ cells/mL) encapsulated in a 1wt% alginate hydrogel cross-linked with Ca^2+^ as previously described to ensure consistent tumorigenesis^60^. Mice were then observed for the development of palpable tumors. One day prior to treatment (day six), tumors were measured, and treatments were randomized and assigned to ensure a comparable tumor load across experimental groups at the start of treatment (40 - 60mm^2^ in tumor area). Seven days post-inoculation, mice were given appropriate treatments of 100μL 1x PBS peritumorally injected for the no treatment control, 100μL of 1x PBS with 2.5μg of cytokine and 5×10^6^ stimulated adoptive T cells retro-orbitally for the intravenous injection, or 100μL of ACTIVATE encapsulating 5×10^6^ stimulated adoptive T cells and 2.5μg of cytokine(s) peritumoral to the tumor. For the synergistic ICB study, 250μg of anti-PD-1 antibody (BioX Cell, Clone RMP1-14) in 100μL of 1x PBS was dosed intraperitoneally once every three days for three doses. For survival studies, tumor progression was assessed three times weekly using digital calipers (Mitutoyo), with tumor area calculated as *length x width*. Mice were euthanized once the total tumor area exceeded 200mm^2^. Body weight was monitored to evaluate short-term toxicity, with mice being euthanized if they experienced a sustained weight loss exceeding 20% from baseline. For flow cytometry studies, mice were euthanized either three days or seven days post-treatment.

### Tumor-draining Lymph Node, Spleen, Tumor, and Gel Sample Preparation for Flow Cytometry

Tumor-draining inguinal lymph nodes and spleen were collected from mice seven days after treatment. Lymph nodes and spleen were mechanically processed into single-cell suspensions using single-frosted microslides (Corning, 2948-75X25) and filtered through 70 μm strainers (Celltreat, 229484). These suspensions were then centrifuged at 300 x g for 5 minutes, resuspended in 1x PBS, and assessed for viability using acridine orange/propidium iodide staining solution (Vitascientific, LGBD10012) on a Luna-FL dual fluorescence cell counter (Logos Biosystems). One million viable cells from each sample were transferred to a 96-well conical-bottom plate (Thermo Scientific, 249570) for staining.

Tumors were similarly excised, weighed, and mechanically processed using surgical scissors. Tumors were mechanically dissociated using a gentleMACS Octo Dissociator (Miltenyi) using the associated mouse Tumor Dissociation Kit (Miltenyi, 130-096-730). Single-cell suspensions were made by passing tumors through a 70 μm cell strainer, centrifuged and resuspended in 1x PBS, and counted as described above. One million viable cells from each sample were transferred to a 96-well conical-bottom plate for staining.

Gels were similarly excised into RPMI and mechanically dissociated using surgical scissors. Samples were shaken at 200 rpm at 37°C for 30 minutes before being pipetted up and down. This was repeated until few gel particulates remained. Samples were then filtered through a 70 μm cell strainer, centrifuged, resuspended, and counted as described above. One million viable cells from each sample were transferred to a 96-well conical bottom plate for staining. Gels from *in vitro* studies were similarly processed.

### Staining Protocol and Flow Cytometry

All samples were initially stained with 200μL of Ghost Dye Violet 510 viability dye (Tonbo Biosciences, 13-0870-T100) on ice for 5 minutes then quenched with 100μL of FACS buffer (1x PBS with 3% heat-inactivated FBS and 1mM EDTA). Samples were centrifuged for 2 minutes at 935 x g. Cells were then incubated on ice with 50μL of anti-mouse CD16/CD32 Fc block (BD, 553142, 1:50 dilution) for 5 minutes before applying 50μL of a cocktail of surface antibodies for 30 minutes on ice. Samples were then centrifuged as before. For intracellular cytokine staining, cells were first stimulated with eBioscience Cell Stimulation Cocktail (Thermo Fisher, 00-4970-93) for 4 to 6 hours at 37°C. Cells were then surface stained as described before being fixed with 200μL of 1x Intracellular Fixation Buffer at room temperature for 30 minutes in the dark. Cells were then pelleted at 500 x g for 5 minutes, washing with 200μL of 1x Permeabilization buffer, and centrifuged at 600 x g for 5 minutes. Intracellular cytokine staining was performed by incubating cells in 100μL of staining solution prepared in 1x Permeabilization Buffer for 30 minutes at room temperature in the dark. Cells were washed with 200μL of 1x Permeabilization Buffer, centrifuged as before, and washed once more before final resuspension in 200μL of FACS buffer. Samples were acquired using an Agilent NovoCyte Penteon Flow Cytometer or a BD LSR II Flow Cytometer as specified at the Stanford Shared FACS facility and analyzed with FlowJo software. For fixation and permeabilization of transcription factors like FoxP3 and IFNγ, the FoxP3/Transcription Factor Staining Buffer Set (Thermo Fisher Scientific, 00-5523-00) were used for fixation and permeabilization according to the manufacturer’s instructions.

LN and spleen full antibody stain included anti-I-A/I-E (MHCII) (1:400 dilution; FITC; BioLegend, 107606), anti-CD45 (1:400 dilution; AF700; BioLegend, 103128), anti-CD161 (NK1.1) (1:100 dilution; BV605; BioLegend, 108715), anti-CD11b (1:100 dilution; BD Biosciences, 563553), anti-F4/80 (1:200 dilution; BV421; BioLegend, 123132), anti-CD4 (1:200 dilution; BUV805; BD Biosciences, 612900), anti-CD8a (1:200 dilution; BV785; BioLegend, 100750), anti-CD3 (1:200 dilution; PerCP/eFluor710; Invitrogen, 46003283), anti-CD19 (1:200 dilution; PE-Cy7; BioLegend, 115520), anti-CD11c (1:200 dilution; PE; BioLegend, 117308), anti-CD90.1 (Thy-1.1) (1:200 dilution; APC-eFluor780; Invitrogen, 47090082), anti-XCR1 (1:200 dilution; AF647; BioLegend, 138213), anti-CD44 (1:80 dilution; BV750; BioLegend, 103079), and anti-CD62L (1:80 dilution; BUV737; BD Biosciences, 612833). *In vivo* hydrogel full antibody stain included those listed above as well as anti-Ly-6G (1:200 dilution; BV711; BioLegend, 127643) and anti-Ly-6C (1:200 dilution; BV570; BioLegend, 128030).

Tumor surface antibody stain included anti-I-A/I-E (MHCII) (1:200 dilution; FITC; BioLegend, 107606), anti-CD45 (1:200 dilution; AF700; BioLegend, 103128), anti-CD161 (NK1.1) (1:50 dilution; BV605; BioLegend, 108715), anti-CD11b (1:100 dilution; BD Biosciences, 563553), anti-F4/80 (1:100 dilution; BV421; BioLegend, 123132), anti-Ly-6G (1:100 dilution; BV711; BioLegend, 127643), anti-Ly-6C (1:100 dilution; BV750; BioLegend, 128030), anti-CD4 (1:100 dilution; BUV805; BD Biosciences, 612900), anti-CD8a (1:100 dilution; BV785; BioLegend, 100750), anti-CD3 (1:100 dilution; PerCP/eFluor710; Invitrogen, 46003283), anti-CD19 (1:100 dilution; PE-Cy7; BioLegend, 115520), anti-CD11c (1:100 dilution; PE; BioLegend, 117308), anti-CD90.1 (Thy-1.1) (1:100 dilution; APC-eFluor780; Invitrogen, 47090082), anti-Tim3 (1:80 dilution; BV711), anti-PD-1 (1:80 dilution; FITC), anti-CD25 (1:80 dilution; BV650), anti-CD90.1 (Thy-1.1) (1:100 dilution; APC-Cy7), anti-CD161 (NK1.1) (1:50 dilution; PE-Cy5), and anti-CD86 (1:100 dilution; PE-Cy5; BioLegend, 105016). Tumor intracellular antibody stain included anti-IFN-γ (1:20 dilution; PE-Cy7; BioLegend, 505826), anti-Granzyme B (1:40 dilution; BV421; BioLegend, 396414), and anti-FoxP3 (1:20 dilution; PE; BioLegend, 126404).

*In vitro* hydrogel full antibody stain for murine adoptive cells included anti-CD279 (PD-1) (1:400 dilution; PE; BioLegend, 135206), anti-CD25 (1:40 dilution; BV421; BioLegend, 101923), anti-CD44 (1:80 dilution; PerCP/Cy5.5; BioLegend, 103032), anti-CD62L (1:100 dilution; AF488; BioLegend, 104420), anti-CD4 (1:800 dilution; APC; BioLegend, 100411), anti-CD11b (1:800 dilution; APC; BioLegend, 101211), anti-CD161 (NK1.1) (1:200 dilution; APC; BioLegend, 108709), and anti-CD19 (1:800 dilution; APC; BioLegend, 159805).

*In vitro* hydrogel full antibody stain for human CAR-T cells included anti-CD4 (1:25 dilution; BUV 395; BD Biosciences, 563552), anti-CD8 (1:25 dilution; BUV805; BD Biosciences, 612890), anti-CD279 (PD-1) (1:100 dilution; PE-Cy7; BioLegend, 329917 and anti-CD25 (1:25 dilution; BV650; BioLegend, 302633).

## Statistical Analysis

For *in vitro* experiments, *n* = 3 was used per timepoint per group, and data are presented as mean +/- standard error of mean (SEM) as specified in the corresponding figure captions. Statistics are ordinary one-way ANOVA with Tukey’s multiple comparisons test run in GraphPad Prism. For *in vivo* studies and flow cytometry, a sample size of *n* = 6 to 10 was used, and mice were randomly assigned to groups based to ensure consistent tumor burden. Data are presented as mean +/- standard error of mean (SEM) as specified in the corresponding figure captions.

Treatment groups for survival curves were determined by log-rank Mantel-Cox test in GraphPad Prism. For flow cytometry studies, comparisons between multiple groups were determined by Kruskal-Wallis test with Dunn’s multiple comparisons test in GraphPad Prism. *P* values less than 0.2 are shown in the text and figures, and all *P* values for the survival curves are in the Supplementary Materials.

## Ethical Statement

All animal procedures were performed according to Stanford APLAC approved protocols.

## Data Availability

All data supporting the results in this study are available within the article and its Supplementary Information. The broad range of raw datasets acquired and analyzed (or any subsets thereof), which would require contextual metadata for reuse, are available from the corresponding author upon reasonable request.

## Acknowledgements

The authors would like to thank all members of the Appel lab for their useful discussion and advice throughout this project. The authors were also grateful to the staff of the BioE/ChemE Animal Facility who cared for the mice. AN is thankful for the generous support of the Paul and Mildred Berg Fellowship. JHK is grateful for the support of the Agilent fellowship. NE and JY were supported by the National Science Foundation Graduate Research Fellowship. SJB would like to thank Stanford’s Office for Postdoctoral Affairs for support through a PRISM Baker Fellowship. KG was supported by a fellowship through the Stanford ChEM-H Chemistry-Biology Interface Training Program (5T32GM139791-04). BSO was supported by Eastman Kodak Fellowship. Flow cytometry was performed on an instrument in the Stanford Shared FACS Facility obtained using NIH S10 Shared Instrument Grant (1S10OD026831-01).

