## Supplementary Information for "A Transient Immunostimulatory Niche Synergizes Adoptive and Endogenous Immunity for Enhanced Tumor Control"

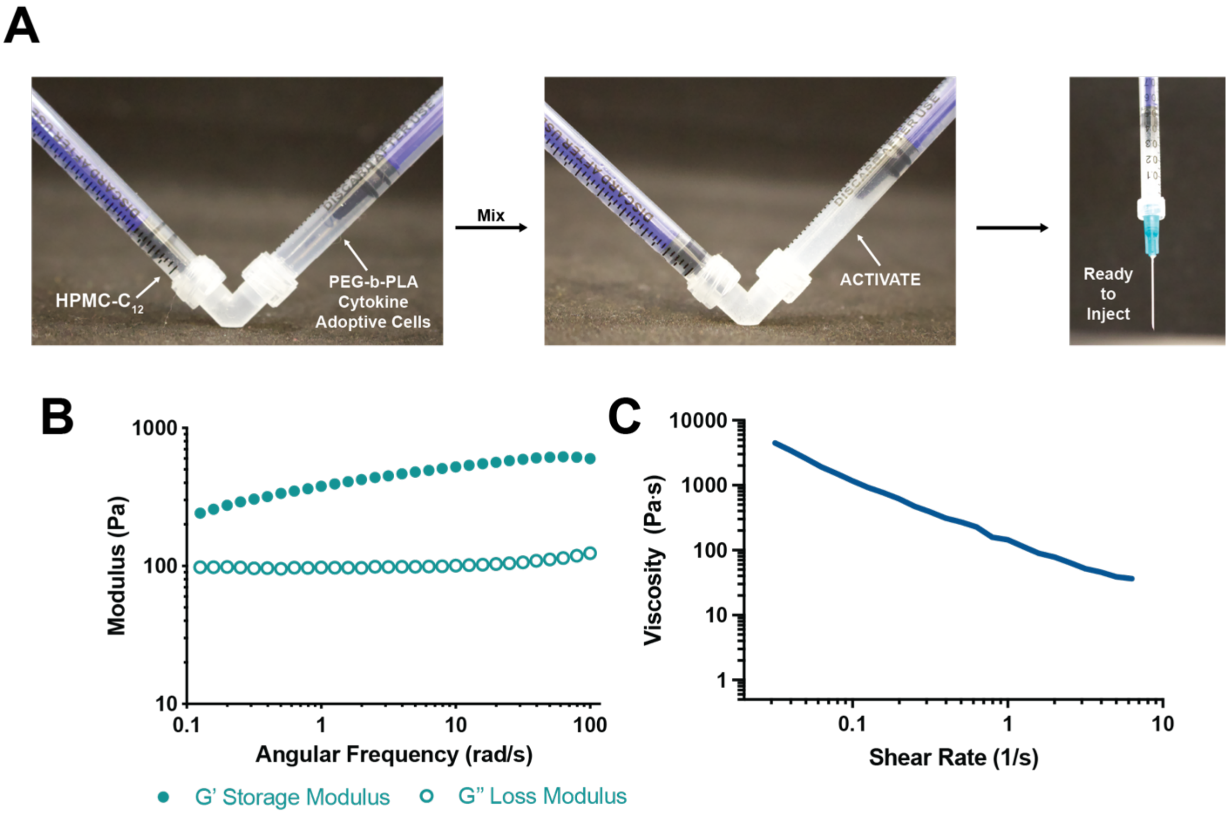


**Figure S1. ACTIVATE Characterization.** (**A**) Facile formulation of ACTIVATE by simple mixing of HPMC-C12 biopolymer solution in one syringe (left) and a mixture of PEG-b-PLA nanoparticles, adoptive cells, and cytokines in the other syringe (right) using a Luer lock elbow mixer. After gentle mixing for 30 seconds, a solid-like PNP hydrogel encapsulating cells homogenously is formed (right syringe). Injection is then performed through a 23-gauge needle. (**B**) Frequency sweep at 1% strain of the 1wt% HPMC-C12 and 5wt% PEG-PLA NPs formulation. (**C**) Flow sweep at high shear rates (representative of injection) for the aforementioned PNP formulation.

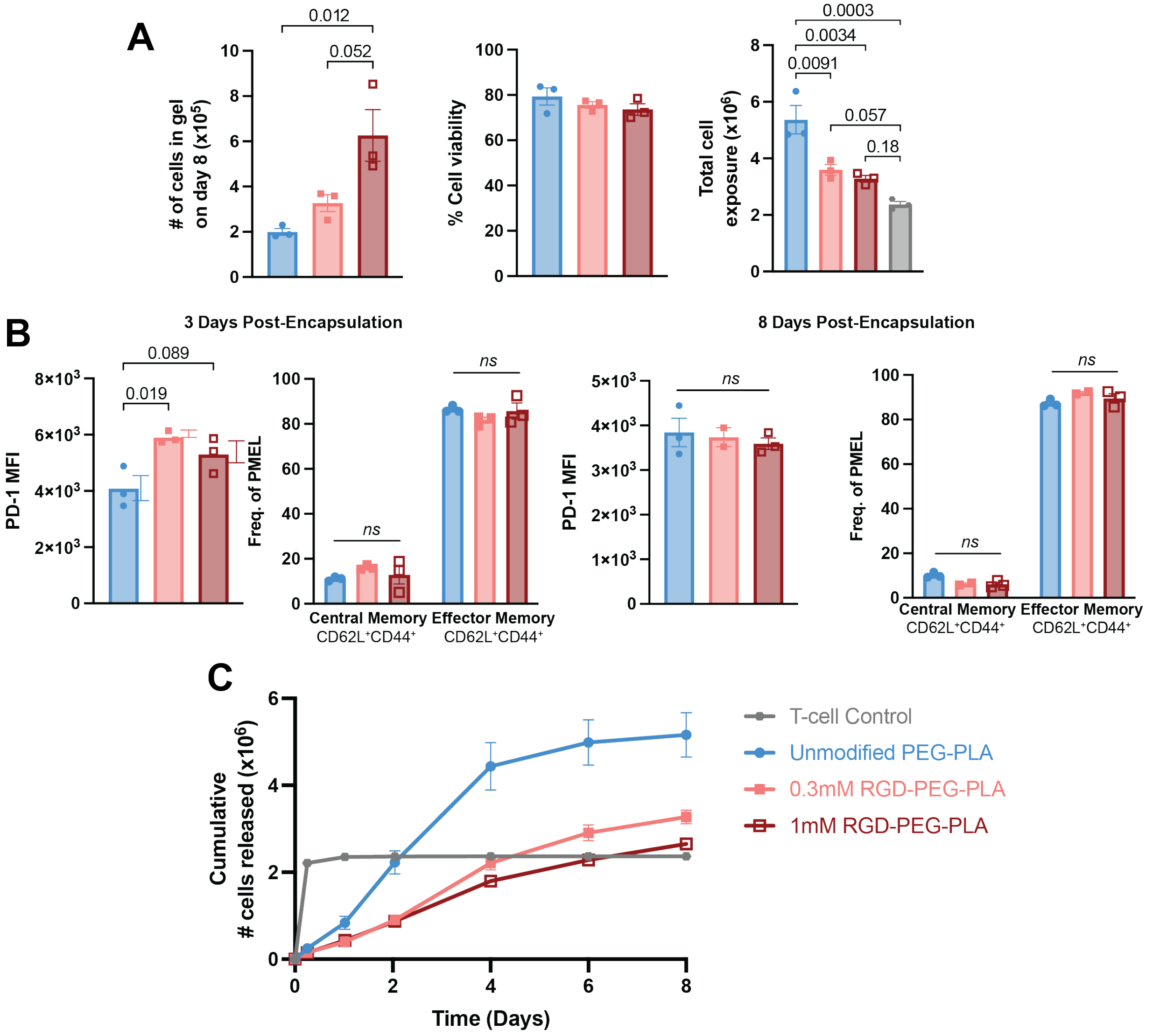


**Figure S2. Characterizing RGD-functionalized PEG-PLA and its effect on T-cell expansion and phenotype.** (**A**) Cumulative number of cells released into the medium over various timepoints of hydrogels encapsulated with 2x106 PMEL cells, 2.5ug of IL-2, and either unmodified PEG-PLA NPs, 0.3mM RGD-modified PEG-PLA NPs, or 0.5mM RGD-modified PEG PLA NPs. (**B**) Total number of cells remaining in various ACTIVATE (left) and cell viability of the cells remaining in the hydrogel (middle) on the last day of the assay (day 8), and total cell exposure (right) determined as the sum of cumulative cells released and total cells remaining in the hydrogel by day 8. (**C**) Quantification of PD-1 MFI of PMEL cells in the gel 3 days after encapsulation (top left) and 8 days post encapsulation (bottom left). Frequency of central memory ($CD62L^+CD44^+$) and effector memory ($CD62L^-CD44^+$) phenotypes for PMEL cells in the hydrogel 3 days after co-encapsulation (top right) and 8 days post-encapsulation (bottom right). Data are representative of *n* = 3 technical replicates per group and presented as mean ± s.e.m. *P* values were determined by ordinary one-way ANOVA followed by Tukey's multiple comparison test using GraphPad PRISM.

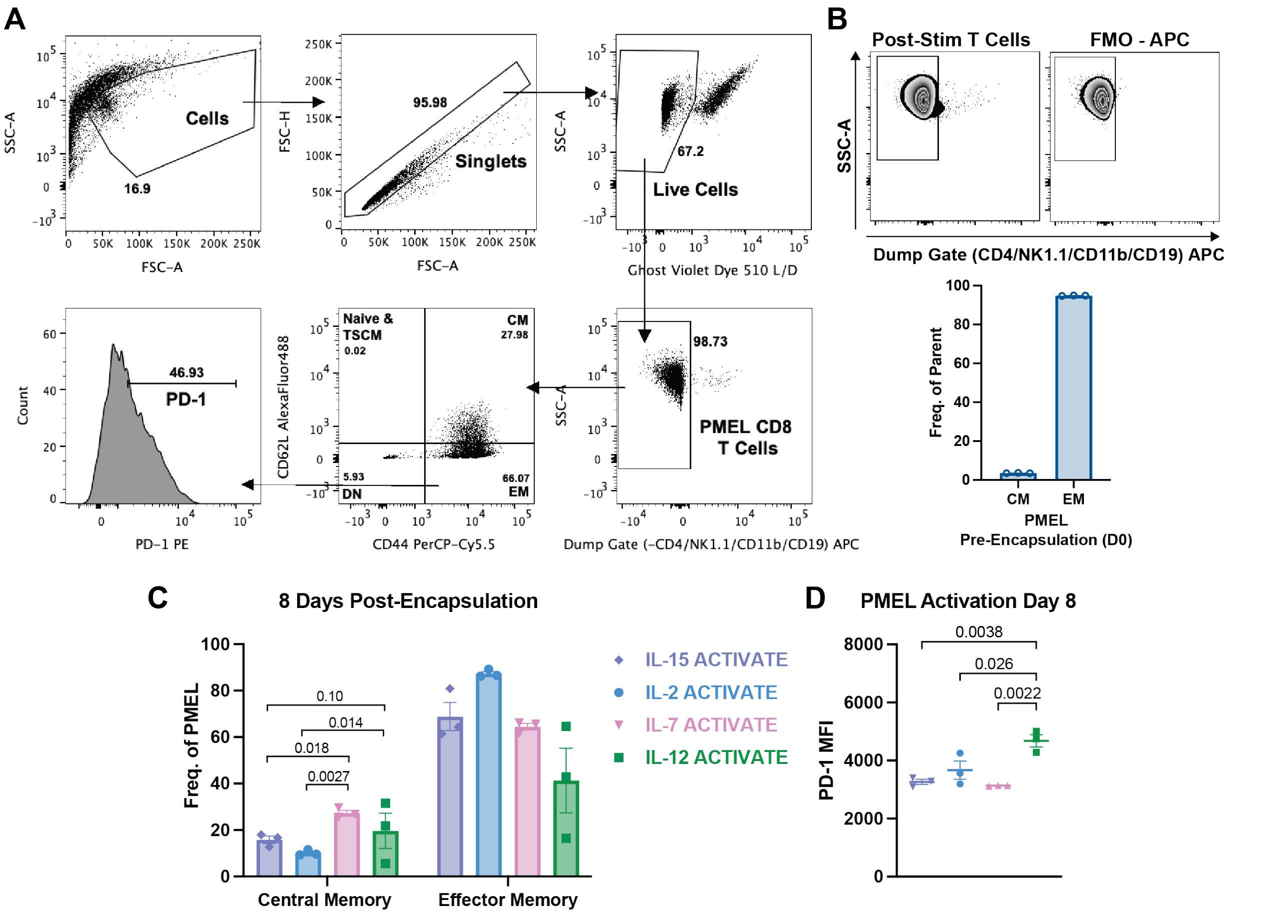


**Figure S3. Cytokines in ACTIVATE influence adoptive T-cell expansion and proliferation.** (**A**) Representative gating strategy for cells in gel harvested and stained either three days or eight days post-encapsulation. (**B**) Representative gating of PMEL cells ten days after splenocyte harvesting, stimulation, and activation in vitro (top). Characterization of the central memory central memory ($CD62L^{+}CD44^{+}$) and effector memory ($CD62L^{-}CD44^{+}$) phenotypes for these cells post-stimulation and prior to encapsulation in ACTIVATE gels. (**C**) Frequency of central memory and effector memory phenotypes for PMEL cells in the hydrogel eight days after encapsulation with various cytokines. (**D**) Quantification of PD-1 MFI of PMEL cells in in the hydrogel eight days after encapsulation with various cytokines. Data are representative of $n$ = 3 technical replicates per group and presented as mean ± s.e.m. $P$ values were determined by ordinary one-way ANOVA followed by Tukey's multiple comparison test using GraphPad PRISM.

**Supplementary Table 1.** Additional P values for treatment groups from Figure 3 survival analysis computed using a log-rank Mantel-Cox test.

**Figure 3B B16-F10**

| Treatment pair | P value |
|---|---|
| NT, IL-12 ACTIVATE | 0.0071 |
| NT, IV + IL-2 | 0.1319 |
| NT, IL-2 ACTIVATE | 0.3092 |
| NT, IL-15 ACTIVATE | 0.014 |
| NT, IL-7 ACTIVATE | 0.8079 |
| IV + IL-2, IL-2 ACTIVATE | 0.5951 |
| IV + IL-2, IL-15 ACTIVATE | 0.1869 |
| IV + IL-2, IL-7 ACTIVATE | 0.4790 |
| IL-2 ACTIVATE, IL-12 ACTIVATE | 0.0014 |
| IL-7 ACTIVATE, IL-12 ACTIVATE | 0.0400 |
| IL-15 ACTIVATE, IL-12 ACTIVATE | <0.0001 |
| IL-2 ACTIVATE, IL-15 ACTIVATE | 0.0588 |
| IL-15 ACTIVATE, IL-7 ACTIVATE | 0.2935 |
| IL-2 ACTIVATE, IL-7 ACTIVATE | 0.6730 |

**Figure 3E E.G7-Ova**

| Treatment pair | P value |
|---|---|
| NT, IL-12 ACTIVATE | <0.0001 |
| NT, IV + IL-2 | 0.0010 |
| NT, IL-2 ACTIVATE | 0.0004 |
| NT, IL-15 ACTIVATE | 0.0010 |
| NT, IL-7 ACTIVATE | 0.0022 |
| IV + IL-2, IL-2 ACTIVATE | 0.1367 |
| IV + IL-2, IL-15 ACTIVATE | 0.2260 |
| IV + IL-2, IL-7 ACTIVATE | 0.8350 |
| IL-2 ACTIVATE, IL-12 ACTIVATE | 0.3185 |
| IL-7 ACTIVATE, IL-12 ACTIVATE | 0.0456 |
| IL-15 ACTIVATE, IL-12 ACTIVATE | 0.0617 |
| IL-2 ACTIVATE, IL-15 ACTIVATE | 0.5758 |
| IL-15 ACTIVATE, IL-7 ACTIVATE | 0.5263 |
| IL-2 ACTIVATE, IL-7 ACTIVATE | 0.2186 |

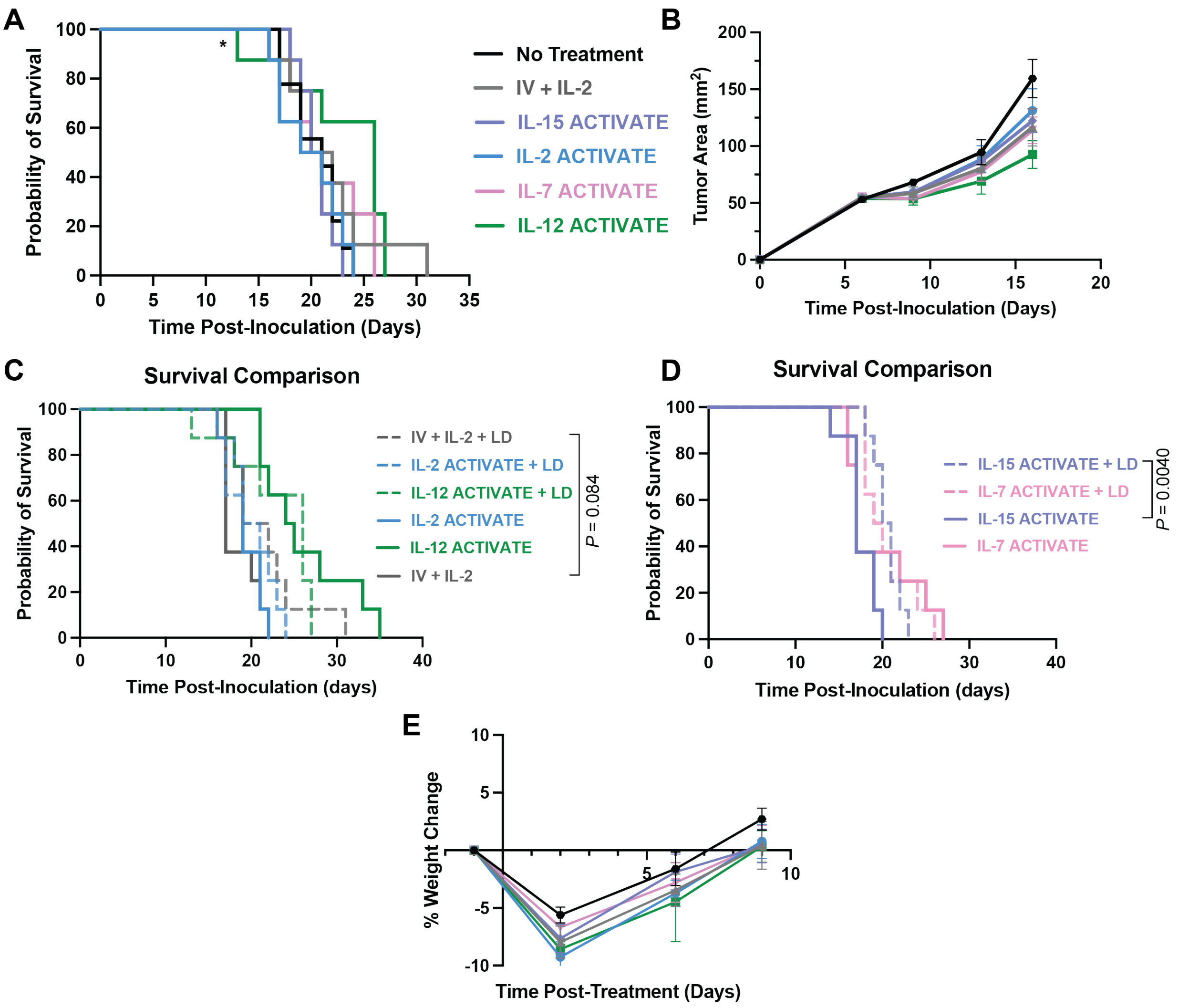


**Figure S4. Lymphodepletion is not needed for T-cell engraftment of ACTIVATE gels and abrogates their anti-tumor efficacy.** C57Bl/6 mice were inoculated (s.c.) with 0.3x10$^{6}$ B16-F10 cells on D0 and treated on D7 with 100uL of ACTIVATE (p.t.) containing 5x10$^{6}$ PMEL T cells and 2.5ug of either IL-2, IL-7, IL-15, or IL-12. I.V. control group received 5x10$^{6}$ PMEL T cells and 2.5ug of IL-2 administered retro-orbitally in a bolus. All mice received lymphodepletion one day prior to treatment (D6). Tumors were measured with digital calipers until tumor burden exceeded euthanasia criteria. (**A**) Survival curve. * indicates one mouse that required euthanasia after sustained weight loss exceeding 20% from baseline. (**B**) Primary tumor growth curve. (**C**) Overall survival comparison of the treatment groups IV + IL-2, IL-2 Gel, and IL-12 Gel with lymphodepletion (dashed lines) and without lymphodepletion (unbroken lines). (**D**) Overall survival comparison of treatment groups IL-7 Gel and IL-15 Gel with lymphodepletion (dashed lines) and without lymphodepletion (unbroken lines). (**E**) % change in mouse body weight over time after treatment. Data are representative of $n$ = 8 and presented as mean ± s.e.m. $P$ values were determined using the log-rank (Mantel-Cox) test using GraphPad PRISM.

**Supplementary Table 2.** Additional P values from Figure S4C and S4D survival analysis computed using a log-rank Mantel-Cox test.

| Treatment pair | P value |
|---|---|
| IL-2 Gel + LD, IL-2 Gel | 0.3693 |
| IL-12 Gel + LD, IL-12 Gel | 0.3441 |
| IL-7 Gel + LD, IL-7 Gel | 0.8567 |

**A** Spleen & Lymph Node Gating Schematic

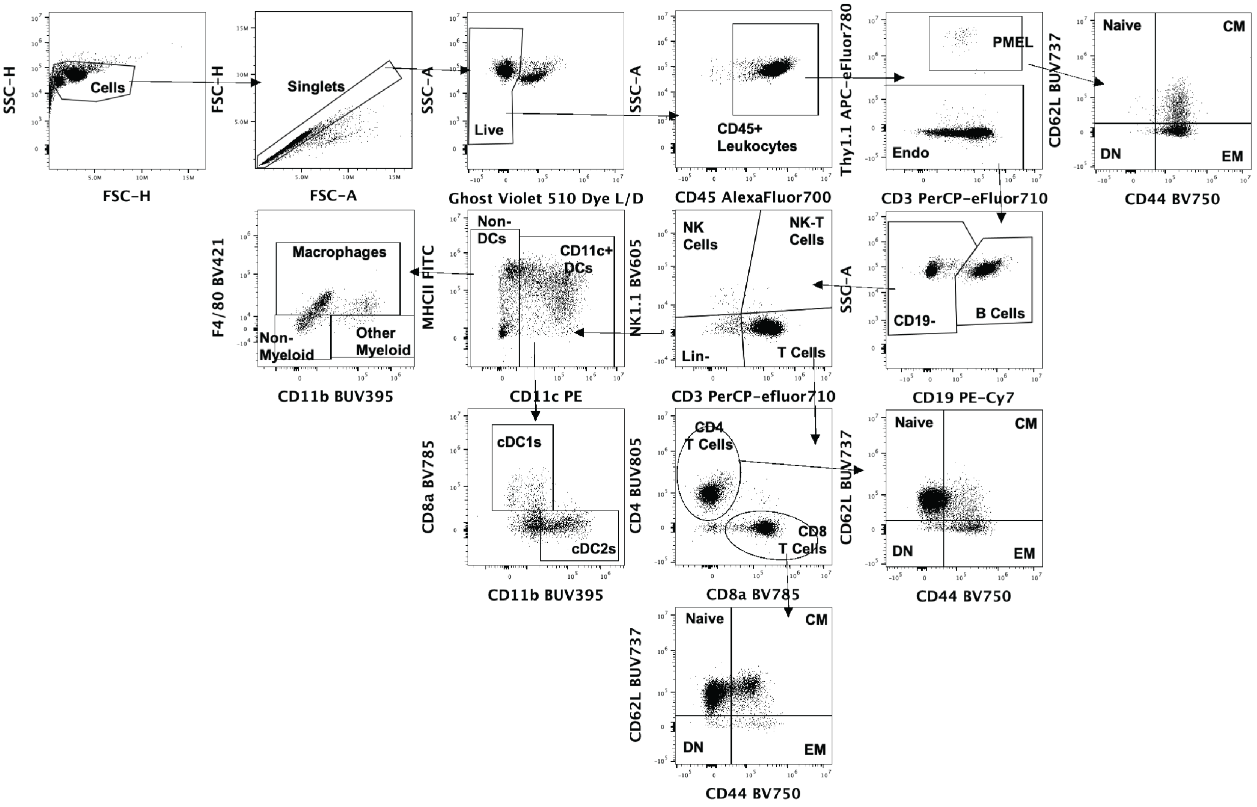


**B** Gated on PMEL or CD8+ Cells from LN

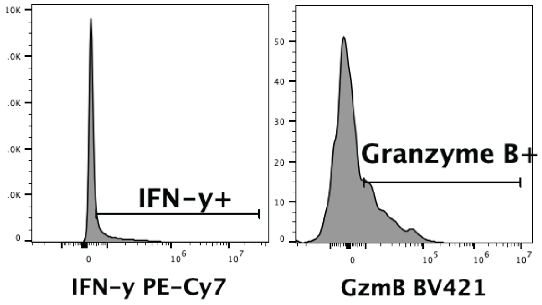


**Figure S5. Representative flow cytometry gating plots for Spleen and Lymph Node samples.**

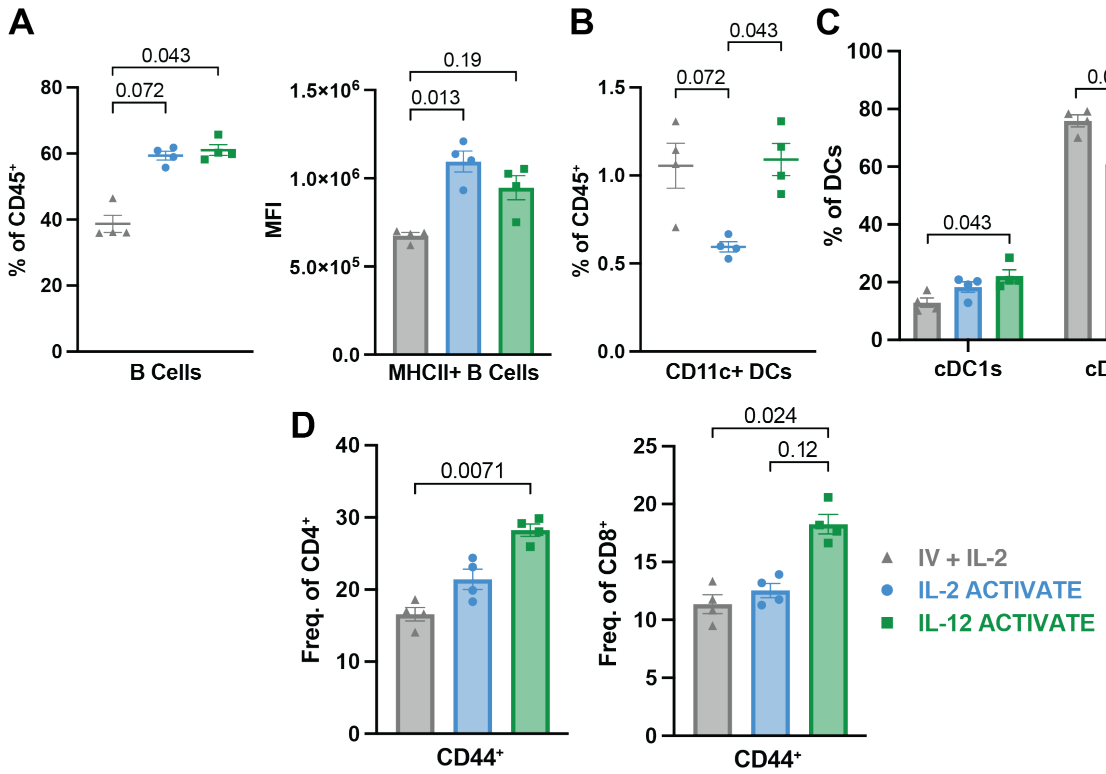


**Figure S6. Characterization of splenocytes from IV or ACTIVATE-treated groups seven days post-treatment.** Experimental design: C57Bl/6 mice were inoculated (s.c.) with 0.3x$10^6$ B16-F10 cells on D0 and treated on D7 with 100uL of ACTIVATE (p.t.) containing 5x$10^6$ PMEL T cells and 2.5ug of either IL-2 or IL-12. 7 days after treatment (D14), spleens were harvested and stained for flow cytometry analysis. (**A**) Frequency of B cells ($CD19^+$) found in the spleen (left) and quantification of $MHCII^+$ B cells (right). (**B**) Frequency of $CD11c^+$ dendritic cells found in the spleen (left). (**C**) Frequency of cDC1s ($CD8^+$) and cDC2s ($CD8^-CD11b^+$) populations from $CD11c^+$ DCs found in the spleen. (**D**) Frequency of $CD44^+$ Helper T cells ($CD4^+$) (left) and $CD44^+$ Cytotoxic T cells ($CD8^+$) (right) found in the spleens. Data are representative of $n$ = 4 for all treatment groups and presented as mean ± s.e.m. $P$ values were determined using Kruskal-Wallis test with Dunn's multiple comparisons test using GraphPad PRISM.

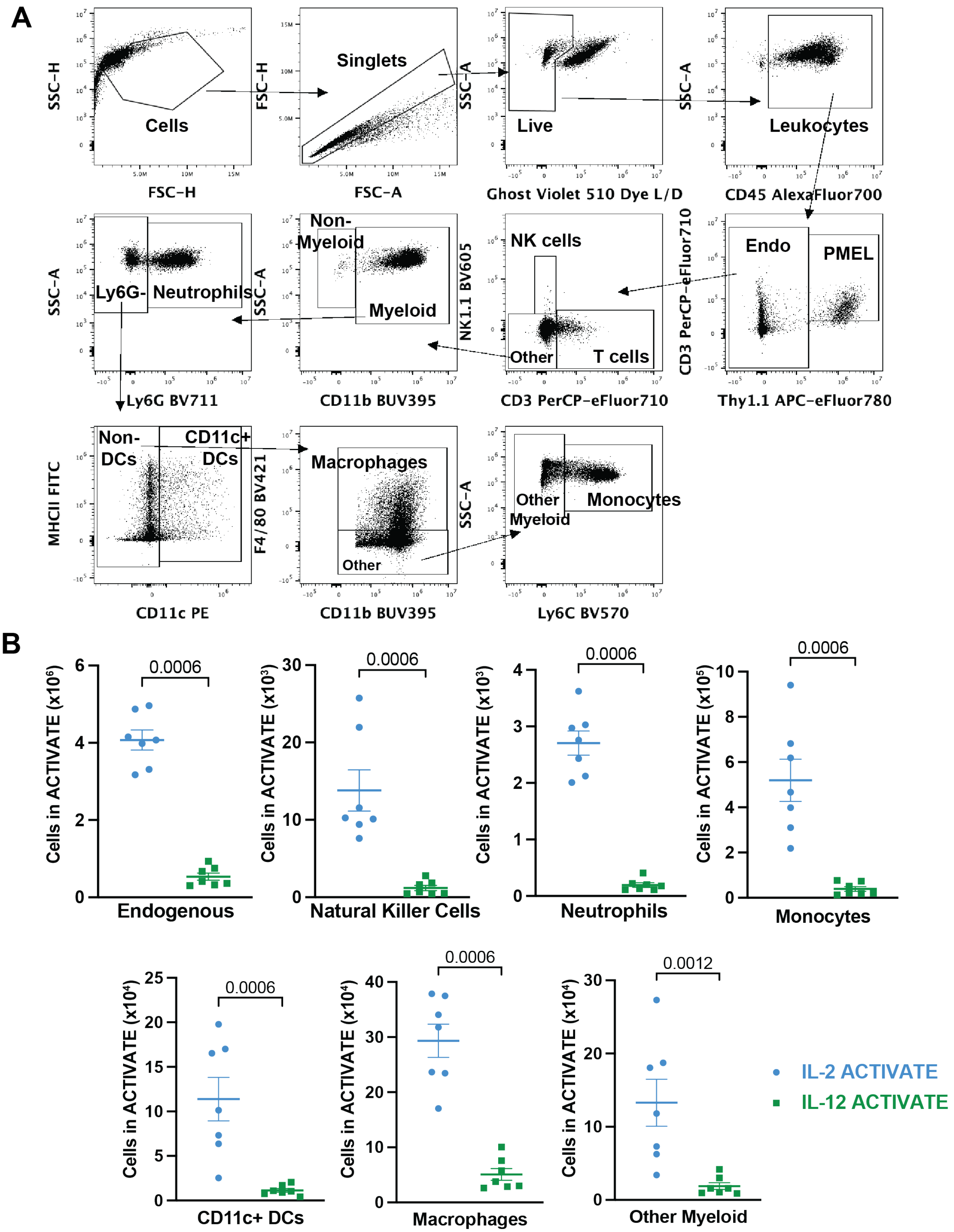


**Figure S7. Flow cytometry characterization of cells harvested from ACTIVATE gels in vivo.** (**A**) Representative flow cytometry gating strategy for cells harvested three days and seven days post-encapsulation from ACTIVATE gels. (**B**) Total cell counts of endogenous cells ($Thy1.1^-$), natural killer cells ($NK1.1^+$), neutrophils ($CD11b^+Ly\text{-}6G^+$), monocytes ($CD11b^+Ly\text{-}6C^+$), $CD11c^+$ DCs, macrophages ($CD11b^+F4/80^+$), and other myeloid ($CD11b^+$) found in the ACTIVATE gels seven days post-encapsulation. Data are representative of $n$ = 7 for both treatment groups and presented as mean ± s.e.m. $P$ values were determined using Mann-Whitney test using GraphPad PRISM.

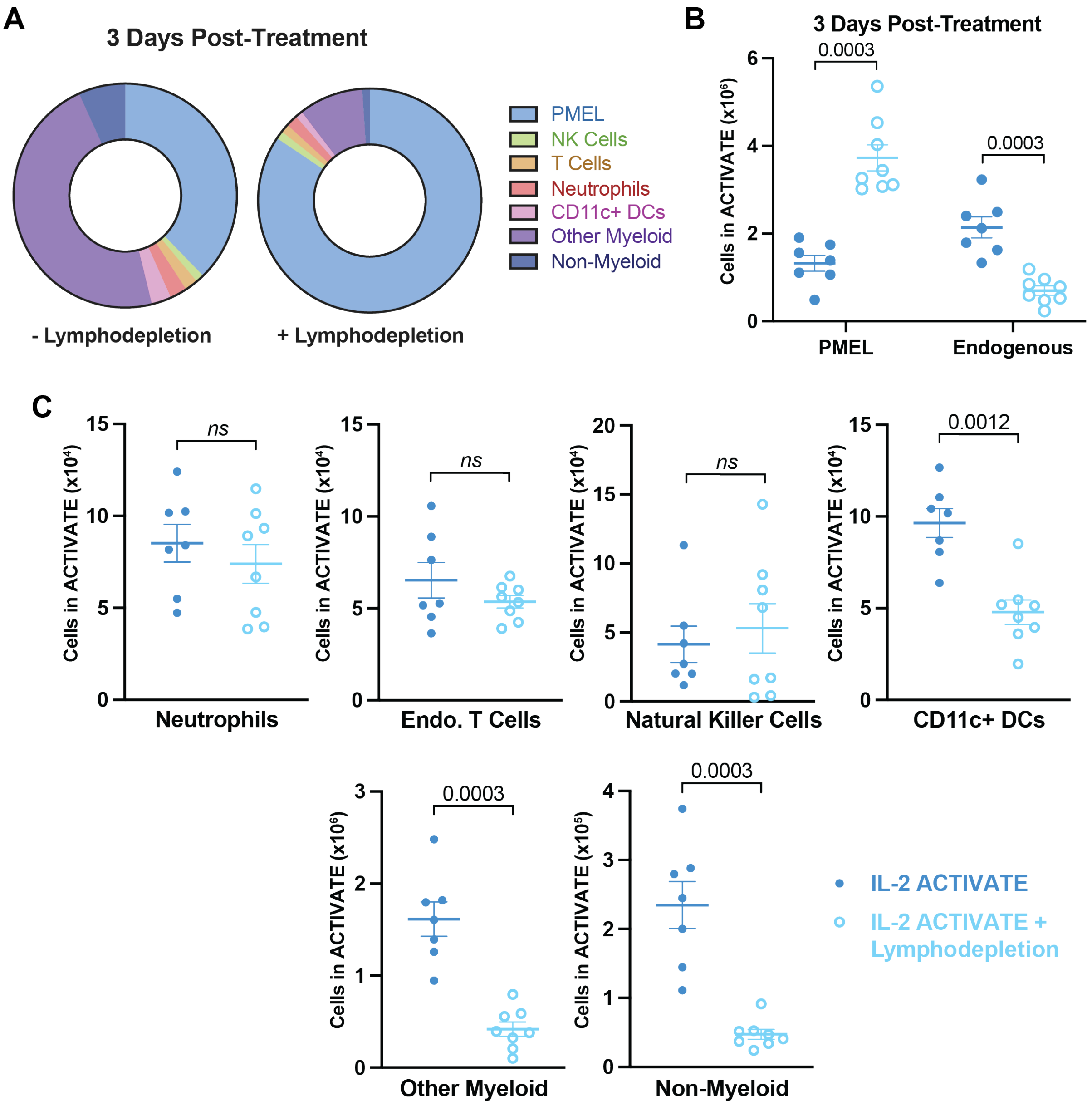


**Figure S8. Comparison of IL-2 ACTIVATE gel with and without lymphodepletion preconditioning regimen.** (**A**) Frequency of PMEL adoptive T cells, natural killer cells, endogenous T cells, neutrophils, $CD11c^+$ dendritic cells, other myeloid, and non-myeloid cells from the $CD45^+$ cell populations found in the IL-2 ACTIVATE gels harvested from mice three days after treatment. (**B**) Total number of PMEL adoptive cells ($Thy1.1^+$) and endogenous cells ($Thy1.1^-$) found in IL-2 ACTIVATE with and without lymphodepletion preconditioning. (**C**) Total number of neutrophils, endogenous T cells, natural killer cells, $CD11c^+$ dendritic cells, other myeloid, and non-myeloid cells found in IL-2 ACTIVATE with and without lymphodepletion three days post-treatment. Data are representative of $n = 7$ per condition and presented as mean ± s.e.m. $P$ values were determined using Mann-Whitney test using GraphPad PRISM.

**Supplementary Table 3.** Additional P values from Figure 7B survival analysis computed using a log-rank Mantel-Cox test.

| Treatment pair | P value |
|---|---|
| NT, IL-2 ACTIVATE | <0.0001 |
| IL-2 ACTIVATE, IL-12 ACTIVATE | 0.0034 |
| NT, IL-2 ACTIVATE | 0.0007 |
| IV + IL-2, IL-2 ACTIVATE | 0.3498 |
| IL-2 ACTIVATE, aPD-1 only | 0.9622 |
| NT, aPD-1 only | 0.0008 |

**Supplementary Table 4.** Additional P values from Figure 7E & 7F compared to Figure 3 survival analysis computed using a log-rank Mantel-Cox test.

**Figure 7E**

| Treatment pair | P value |
|---|---|
| NT, IL-12/IL-7 ACTIVATE | <0.0001 |
| NT, IL-12/IL-2 ACTIVATE | <0.0001 |
| IL-2 ACTIVATE, IL-12/IL-2 ACTIVATE | <0.0001 |
| IL-7 ACTIVATE, IL-12/IL-7 ACTIVATE | <0.0001 |
| IL-12 ACTIVATE, IL-12/IL-7 ACTIVATE | 0.0527 |
| IL-12 ACTIVATE, IL-12/IL-2 ACTIVATE | 0.0039 |

**Figure 7F**

| Treatment pair | P value |
|---|---|
| NT, IL-12/IL-7 ACTIVATE | <0.0001 |
| NT, IL-12/IL-2 ACTIVATE | <0.0001 |
| IL-2 ACTIVATE, IL-12/IL-2 ACTIVATE | 0.0175 |
| IL-7 ACTIVATE, IL-12/IL-7 ACTIVATE | 0.0023 |
| IL-12 ACTIVATE, IL-12/IL-7 ACTIVATE | 0.0727 |
| IL-12 ACTIVATE, IL-12/IL-2 ACTIVATE | 0.1228 |

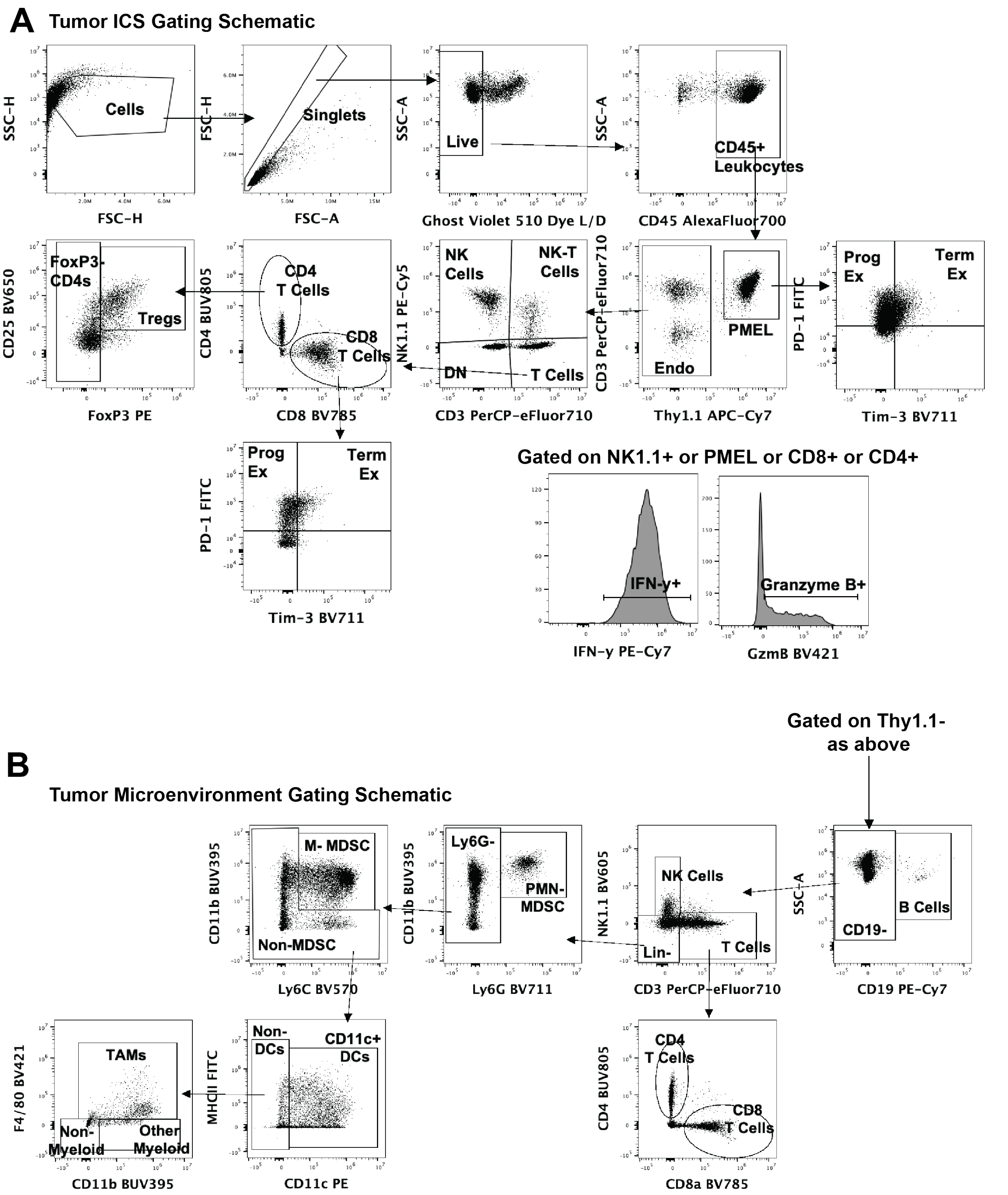


**Figure S9. Representative flow cytometry gating strategies for tumor intracellular cytokine staining (A) and tumor microenvironment staining (B).**

### Supplementary Discussion: Calculation of mIL-2 dose equivalence in humans.

Due to the systemic toxicity associated with cytokines, the dosage in our studies is converted from available literature for the maximum tolerated subcutaneous dose of IL-2 in humans. We chose to focus specifically on IL-2 because of 1) its prevalence in usage alongside PMEL adoptive T cell studies in literature (Ref 39) and 2) because it is the only cytokine immunotherapy currently approved by the FDA (Proleukin, otherwise known as Aldesleukin) for metastatic renal cell carcinoma (RCC) and metastatic melanoma) (Ref 51).

Proleukin is intravenously administered at 600,000 IU/kg three times a day every 8 hours for 14 to 20 doses depending on treatment of metastatic RCC or metastatic melanoma (Ref 52). We chose to scale to recombinant human IL-2 from R&D Systems (202-IL) due to the international unit conversion availability, which is $9.1x10^3$ IU per 1 ug. Taking a single day's worth of Proleukin at 1.8 MIU/kg, we can convert that to ug of IL-2 by dividing by 9,100 IU/ug, which gives 197.8 ug/kg. Assuming mice are approximately 0.02 kg in weight, this gives a dose of 3.96 ug/mouse.

However, in another study, the maximum tolerated subcutaneous dose of IL-2 for one day in humans was determined to be 12 MIU/$m^2$ (Ref 53). Scaling this dosage to mouse by dividing 12 MIU/$m^2$ by $9.1x10^3$ IU/ug, we get 1,318.68 ug/$m^2$. Scaling this dose to mice by dividing by 12.3 (Ref 54) gives 107.25 ug/kg. Again, assuming mice are approximately 0.02 kg in weight, this gives 2.15 ug per day in one dose in a mouse.

We chose to proceed with a dosage of 2.5 ug for IL-2 and each subsequent cytokine used throughout this study, which is lower than the maximum tolerated dose of intravenously delivered IL-2 but slightly higher than the MTD of subcutaneously delivered IL-2.
